# Reconstructing the human genetic history of mainland Southeast Asia: insights from genome-wide data from Thailand and Laos

**DOI:** 10.1101/2020.12.24.424294

**Authors:** Wibhu Kutanan, Dang Liu, Jatupol Kampuansai, Metawee Srikummool, Suparat Srithawong, Rasmi Shoocongdej, Sukrit Sangkhano, Sukhum Ruangchai, Pittayawat Pittayaporn, Leonardo Arias, Mark Stoneking

## Abstract

Thailand and Laos, located in the center of Mainland Southeast Asia (MSEA), harbor diverse ethnolinguistic groups encompassing all five language families of MSEA: Tai-Kadai (TK), Austroasiatic (AA), Sino-Tibetan (ST), Hmong-Mien (HM) and Austronesian (AN). Previous genetic studies of Thai/Lao populations have focused almost exclusively on uniparental markers and there is a paucity of genome-wide studies. We therefore generated genome-wide SNP data for 33 ethnolinguistic groups, belonging to the five MSEA language families from Thailand and Laos, and analysed these together with data from modern Asian populations and SEA ancient samples. Overall, we find genetic structure according to language family, albeit with heterogeneity in the AA-, HM- and ST-speaking groups, and in the hill tribes, that reflects both population interactions and genetic drift. For the TK speaking groups, we find localized genetic structure that is driven by different levels of interaction with other groups in the same geographic region. Several Thai groups exhibit admixture from South Asia, which we date to ∼600-1000 years ago, corresponding to a time of intensive international trade networks that had a major cultural impact on Thailand. An AN group from Southern Thailand shows both South Asian admixture as well as overall affinities with AA-speaking groups in the region, suggesting an impact of cultural diffusion. Overall, we provide the first detailed insights into the genetic profiles of Thai/Lao ethnolinguistic groups, which should be helpful for reconstructing human genetic history in MSEA and selecting populations for participation in ongoing whole genome sequence and biomedical studies.

## Introduction

Mainland Southeast Asia (MSEA), consisting of Myanmar, Cambodia, Vietnam, western Malaysia, Laos, and Thailand, is a region of enormous diversity, with a population of ∼263 million people speaking ∼229 languages belonging to 5 major language families: Tai-Kadai (TK), Austroasiatic (AA), Sino-Tibetan (ST), Hmong-Mien (HM), and Austronesian (AN) (Eberhard, Simons and Fennig, 2020). Thailand and Laos are in the center of MSEA, and are characterised by a diverse landscape involving highlands and lowlands, long coastlines, and many rivers. North-vs.-south movements are facilitated by several rivers, including the Mekong, Chao Phraya, and Salaween which are considered to be a key factor for population movement from southern China and upper MSEA to lower MSEA. In addition, the Malay Peninsula to the south acts as a cross-road, facilitating east-vs.-west movement by sea and by the narrow width of the Kra Isthmus (the narrowest part of the Malay Peninsula).

The geographic heterogeneity of Thailand and Laos is reflected in the ethnolinguistic diversity of the region. There are ∼68.6 million people in Thailand and ∼6.8 million in Laos, speaking ∼159 languages belonging to all five major MSEA language families (Eberhard, Simons and Fennig, 2020). TK languages are widespread in southern China and MSEA, and are quite prevalent in present-day Thailand, and Laos, spoken by 89.4% of Thais and 65.7% of Laotians. The major TK speaking groups in northern, northeastern, central and southern Thailand are known as Khonmueang, Lao Isan, Central Thai, and Southern Thai or Khon Tai, respectively (Eberhard, Simons and Fennig, 2020). AA languages are next in predominance, spoken by 4.0% of Thais and 26.2% of Laotians. In addition, this area is also inhabited by historical migrants who speak ST, HM, and AN languages (frequencies of 3.2%, 0.2%, and 2.8%, respectively, in Thailand; and 2.9%, 4.7%, and 0% in Laos) (Eberhard, Simons and Fennig, 2020). The AA, HM, and ST languages are spoken mainly by highlanders (the hill tribes) in northern and western Thailand, and in midland and upland regions in Laos, although AA languages are also spoken by some lowland groups, e.g. the Mon. AN-speaking groups, such as the Thai Malay (SouthernThai_AN), are distributed in the Southern Provinces of Thailand, bordering with Malaysia.

Archaeological records document a long history of human occupation of the area, with modern human remains dated to 46-63 thousand years ago (kya) in northern Laos (Demeter et al., 2012). In addition, cultural remains of SEA hunter-gatherers (e.g. flake stone tools of the Hòabìnhian culture) have been found in northern Thailand dating to 35-40 kya (Shoocongdej, 2006), and in southern Thailand dating to 27-38 kya (Anderson, 1990). The transition from a hunter-gatherer tradition to a Neolithic agricultural lifestyle occurs ∼4 kya all across Thailand and Laos (Higham and Thodsarat, 2012; Higham, 2014); agriculture in MSEA probably has its origins in the valley of the Yangtze River in China (Higham and Thodsarat, 2012), and ancient DNA evidence indicates that present-day AA speaking groups in MSEA are most closely related to Neolithic agricultural communities (McColl et al., 2018; Lipson et al., 2018).

However, the common languages shared by Thais and Laotians are TK languages, not AA languages. The origin of the TK languages is thought to be in what is now southern or southeastern China, and they probably spread to MSEA during the Iron Age (Pittayaporn, 2014). Whether the spread of TK languages occurred via demic diffusion (an expansion of people that brought both their genes and their language) or cultural diffusion (language spread with at most minor movement of people) has been debated (Nakbunlung, 1994; Sangvichien, 1966; Pittayaporn, 2014). Previous genetic studies of uniparental lineages have generally supported demic diffusion for the maternal side but cultural diffusion from the AA people for the paternal side for major Thai/Lao TK groups (Kutanan et al., 2017, 2018b, 2019). Archaeological evidence suggests other population contacts in the region, e.g. objects from India that appear during the late Bronze Age and Iron Age and involve the AA-speaking Khmer and Mon (Higham and Thodsarat, 2012; Higham, 2014). Moreover, the HM- and ST-speaking hill tribes in the mountainous areas of northern Thailand, northern Myanmar, northern Laos and southern China migrated to the region during historical times, ∼200 years ago (ya) (Schliesinger, 2000; Penth and Forbes, 2004). Taken together, the archaeological and linguistic evidence suggests a complex population structure and history of the ethnolinguistic groups of Thailand and Laos.

This population structure and history remains largely unexplored by genetic studies, which have almost exclusively analyzed autosomal short tandem repeat (STR) loci, and mitochondrial DNA (mtDNA) and male specific Y chromosome (MSY) sequences. These studies revealed the relative genetic heterogeneity of the AA groups and homogeneity of TK groups (Kampuansai et al., 2017, 2020; Kutanan et al., 2014, 2017, 2019; Srithawong et al., 2015, 2020) and contrasting male and female genetic histories in the region, especially for the matrilocal vs. patrilocal hill tribes (Oota et al., 2001; Besaggio et al., 2007; Kutanan et al., 2018a, 2019, 2020). While genome-wide data provide much richer insights into population structure and genetic history, previous genome-wide studies of Thai/Lao populations are either primarily from northern populations (HUGO Pan-Asian SNP Consortium, 2009; Xu et al., 2010; Lipson et al., 2018) or do not provide any information on ethnolinguistic background (Wangkumhang et al., 2013; Lazaridis et al., 2014). Therefore, we here generated genome-wide SNP data for 452 individuals from 33 ethnolinguistic groups from Thailand and Laos, including two southern Thai groups that have not been involved in any previous genetic studies, speaking languages that encompass all five language families in MSEA. We analysed the allele and haplotype sharing within and between the Thai/Lao groups, and compared them with both modern Asian populations and nearby SEA ancient samples. Our results provide several new insights into the genetic prehistory of MSEA through the lens of populations from Thailand and Laos.

## Results

### Genetic structure and genetic relationships within and between Thai/Lao and other Asian populations

#### Principal Components Analysis (PCA)

We generated genome-wide SNP data for 452 individuals from 32 populations from Thailand and one population from Laos; when combined with previously published data from three Thai populations (Lipson et al., 2018; Lazaridis et al., 2014), there are 482 Thai/Lao samples belonging to 36 populations (Figure 1). We also merged our data with data from modern Asian populations generated on the same platform and SEA ancient samples (Supplementary Table 1; Supplementary Figure 1). We began with PCA to investigate the overall population structure of the merged dataset and identify any outliers (Supplementary Figure 2). After outliers were removed, PC1 separates South Asian (SA) from East Asian (EA) groups, with the Kharia (#44), Onge (#45), and Uygur (#65) located in between (Figure 2A; Supplementary Figure 3). PC2 separates Northeast Asian (NEA) groups from SEA groups. With respect to the major MSEA linguistic groups, ST and HM groups are generally separated from the AA, TK, and AN groups on PC2, while the latter three overlap one another. Exceptionally, the Karen speaking ST groups (Karen_ST; #7-9) also overlap the AA, TK, and AN groups (Figure 2B), while the ST-speaking Lahu from Thailand (#6) and China (#56) and the HM-speaking IuMien (#3) are grouped with the AA-speaking Kinh (#52) and close to the northern Thai TK groups (N_TK; #21-26). Strikingly, four Thai groups from this study, i.e. the AA-speaking Mon (#20), AN-speaking SouthernThai_AN (#4), and TK-speaking CentralThai (#34) and SouthernThai_TK (#35), as well as the previously-published Thai-HO (#36; this population is from the Human Origins dataset of Lazaridis et al., 2014, with no further details available), Mamanwa (#46) and Cambodian (#51), all show additional affinity toward the SA populations (Figure 2A-B).

**Figure 1.**
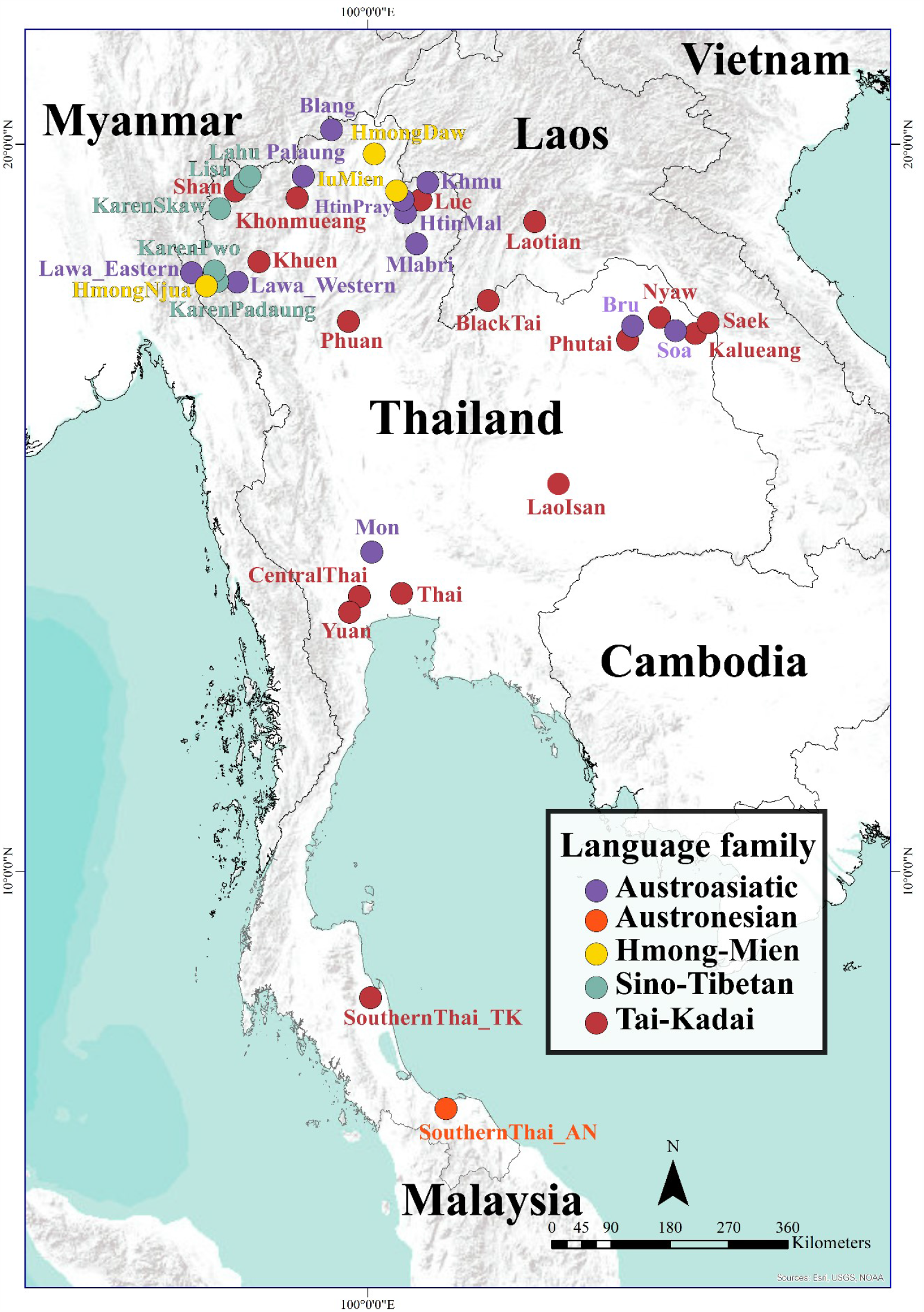
Map showing the location of the 36 Thai/Lao ethnolinguistic groups analyzed in this study, color-coded according to language family.

**Figure 2.**
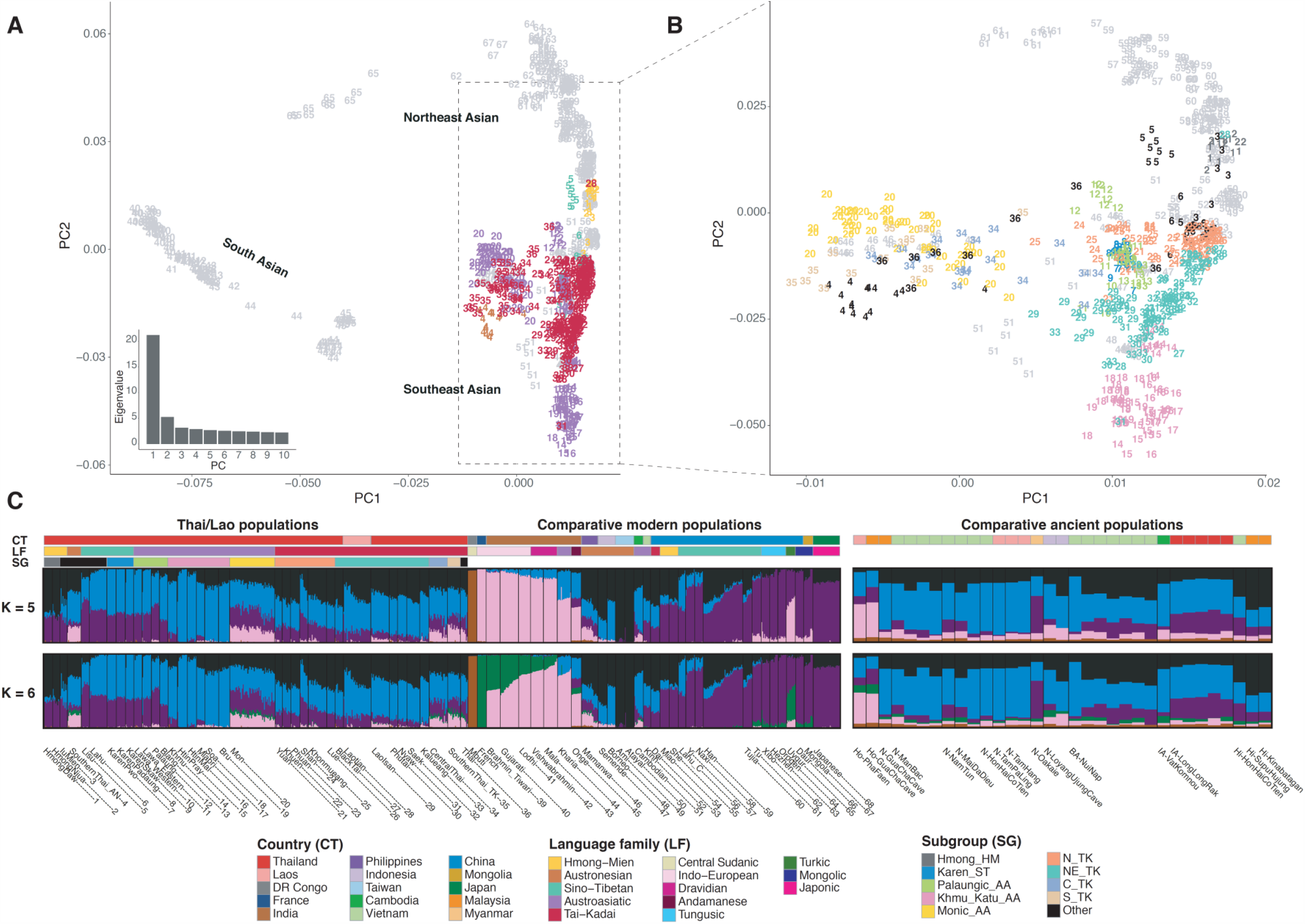
Population structure analyses. (A) Plot of PC1 vs. PC2 for the SNP data for individuals from South Asia, Northeast Asia and Southeast Asia. Individuals are numbered according to population, as indicated in Supplementary Table 1 and in the population labels in panel (C). Thai/Lao groups are colored by language family according to the key at the bottom of panel (C) while other groups are in grey (see Supplementary Figure 3 for the same PC plot with all samples colored by country and by language family). The eigenvalues from PC1 to PC10 are shown on the bottom left side. (B) Plot focusing on Southeast Asian and Chinese populations speaking AA, AN, HM, ST, and TK languages, zoomed-in from (A). Thai/Lao groups are colored according to subgroup while other groups are in grey. (C) ADMIXTURE results for *K* = 5 and *K*= 6. Each individual is represented by a bar, which is partitioned into *K* colored segments that represent the individual’s estimated membership fractions in each of the *K* ancestry components. Populations are separated by black lines for modern populations and excavation sites and time periods are separated by black lines for ancient samples. The three colored bars at the top of the plot indicate the country (top), language family (middle) and subgroup (bottom) for each sample, according to the key at the bottom. The PCA analysis was performed on the pruned dataset of 842 individuals and 153,191 SNPs, while the ADMIXTURE analysis was performed on the pruned dataset of 895 individuals (including 10 Mbuti, 10 French, and 33 ancient individuals) and 158,772 SNPs; the highly drifted modern populations (Onge, Mlabri, and Mamanwa) and ancient samples were projected in ADMIXTURE analyses (see PCA with ancient samples projected in Supplementary Figure 3).

Based on the PCA (Figure 2B), the Thai AA speaking groups can be roughly divided into three groups: Palaungic_AA (Lawa_Western, Lawa_Eastern, Palaung and Blang; #10-13); Khmu_Katu_AA (Khmu, HtinPray, HtinMal, Mlabri, Soa and Bru; #14-19); and Monic_AA (Mon; #20). This grouping is also consistent with their linguistic classification, e.g. Palaungic, Khmuic and Katuic, and Monic (Diffloth, 2005; Sidwell, 2014). The TK groups from different geographic regions in Thailand show different relationships; the N_TK groups are close to the Palaungic_AA groups, AA-speaking Kinh, AN groups from Taiwan (#49-50) and the Philippines (#46), while the northeastern Thai TK groups (NE_TK; Black Tai, Lao Isan, Phutai, Nyaw, Saek and Kalueang; #27 and #29-33) are close to the Khmu_Katu_AA groups. The TK speaking Laotian (#28) are grouped with the NE_TK groups. The central and southern Thai TK groups (C_TK and S_TK; CentralThai and SouthernThai_TK; #34 and #35) and Thai-HO (#36) are close to the Monic_AA groups. Interestingly, the AN-speaking group from Thailand (SouthernThai_AN; #4), is not close to the AN groups from Taiwan (Ami and Atayal) or Indonesia (Semende and Borneo; #47-48), but rather they are near the AN-speaking Negrito group Mamanwa (#46) from the Philippines, and the Monic_AA, C_TK and S_TK groups. Notably, we found two distinct clusters of Mamanwa groups, one is close to N_TK groups, while the other is placed with those groups toward the SA side.

When ancient samples are included in the PCA (Supplementary Figure 3), the two Hòabìnhian samples (#69-70) are projected close to the Onge, while most of the Neolithic samples (#71-79) fall with the AA and AN groups. However, the N-Oakaie sample (#78) from Myanmar is closer to ST and HM groups. Most of the Bronze/Iron Ages samples (#80-82) cluster with the TK and AA samples except for the BA-NuiNap samples (#80) from Vietnam, which are close to the Neolithic samples.

#### ADMIXTURE analysis

We then performed ADMIXTURE analysis to investigate population structure. The lowest cross validation error occurred at *K* = 5 and *K* = 6 (Supplementary Figure 4); corresponding results are shown in Figure 2C. For *K* = 5, there is a brown component associated with Mbuti, a pink component appearing in French and Indian groups, a purple component enriched in NEA groups, a black component dominant in AN-speaking Ami and Atayal from Taiwan, and a blue component enriched in Khmu_Katu_AA groups from Thailand. Most of the Thai/Lao TK-speaking groups show two major sources (black and blue) with the purple component as a minor source, except that the C_TK and S_TK groups and Thai-HO have a substantial fraction of the pink component, as do the Monic_AA and Southern Thai_AN. This indication of potential relatedness with SA groups is consistent with the PCA results (Figure 2A-2B). Also in accordance with the PCA results, the AA-speaking groups can be categorized into 3 groups: the Palaungic_AA group exhibits two major sources (blue and purple) with the black component as a minor source; the Monic_AA group possesses the pink component; and the Khmu_Katu_AA group has a reduced frequency of the purple component.

With respect to the ancient samples at *K* = 5 (Figure 2C), the Hòabìnhian samples show a major pink component with minor blue and purple components, while all of the Neolithic samples exhibit a major blue component with minor black, pink, and purple components, except that the purple component is enriched in the N-Oakaie sample from Myanmar, and reduced/lacked in the N-GuaChaCave samples from Malaysia and the N-TamPaLing and N-TamHang samples from Laos. The purple component is also enriched in the Iron Age samples IA-LongLongRak from Thailand. The black component is substantially increased in the Bronze Age and historical samples, such as the BA-NuiNap and Hi-HonHaiCoTien samples from Vietnam and the Hi-SupuHujung and Hi-Kinabatagan samples from Malaysia (a similar pattern is seen in the Thai/Lao TK groups).

At *K* = 6, there appears a green component that separates French from South Asian populations (Figure 2C). This green component substantially reduces the pink component in the NEA groups, but has a negligible effect on the SA-related Thai groups. Although increasing *K* values are associated with higher cross-validation errors, the additional new components reveal additional population structure (Supplementary Figure 5). At *K* = 7, 8 and 9, the Lahu from Thailand and China, the Hmong_HM, and Karen_ST groups from Thailand are enriched for their own sources, respectively. At *K* = 11, the Soa and Bru (Katuic speaking populations of the Khmu_Katu_AA group) stand out with a light brown component, and in accordance with the PCA results, the different TK-speaking groups can be distinguished: the blue component is now enriched mostly in the N_TK group, the additional light brown component is enriched in the NE_TK group, and the C_TK and S_TK group possess the additional pink component as mentioned previously.

#### Outgroup f3

To further analyse population relationships based on allele sharing, we calculated outgroup *f3-* statistics of the form *f3*(X, Y; outgroup) that measure the shared drift between populations X and Y since their divergence from the outgroup (Mbuti). Higher outgroup *f3* values indicate more shared drift between populations. The SouthernThai_AN, Monic_AA, C_TK, and S_TK groups and Thai-HO exhibit the lowest *f3*-values with other populations/ancient samples and also with each other (Figure 3), while the HM speaking populations show the strongest sharing with each other. TK populations exhibit close genetic affinity with each other, except for the C_TK, S_TK, and Thai-HO groups, and also share alleles with the HM speaking populations, consistent with results of the ADMIXTURE analysis at *K* = 8 (Supplementary Figure 5).

**Figure 3.**
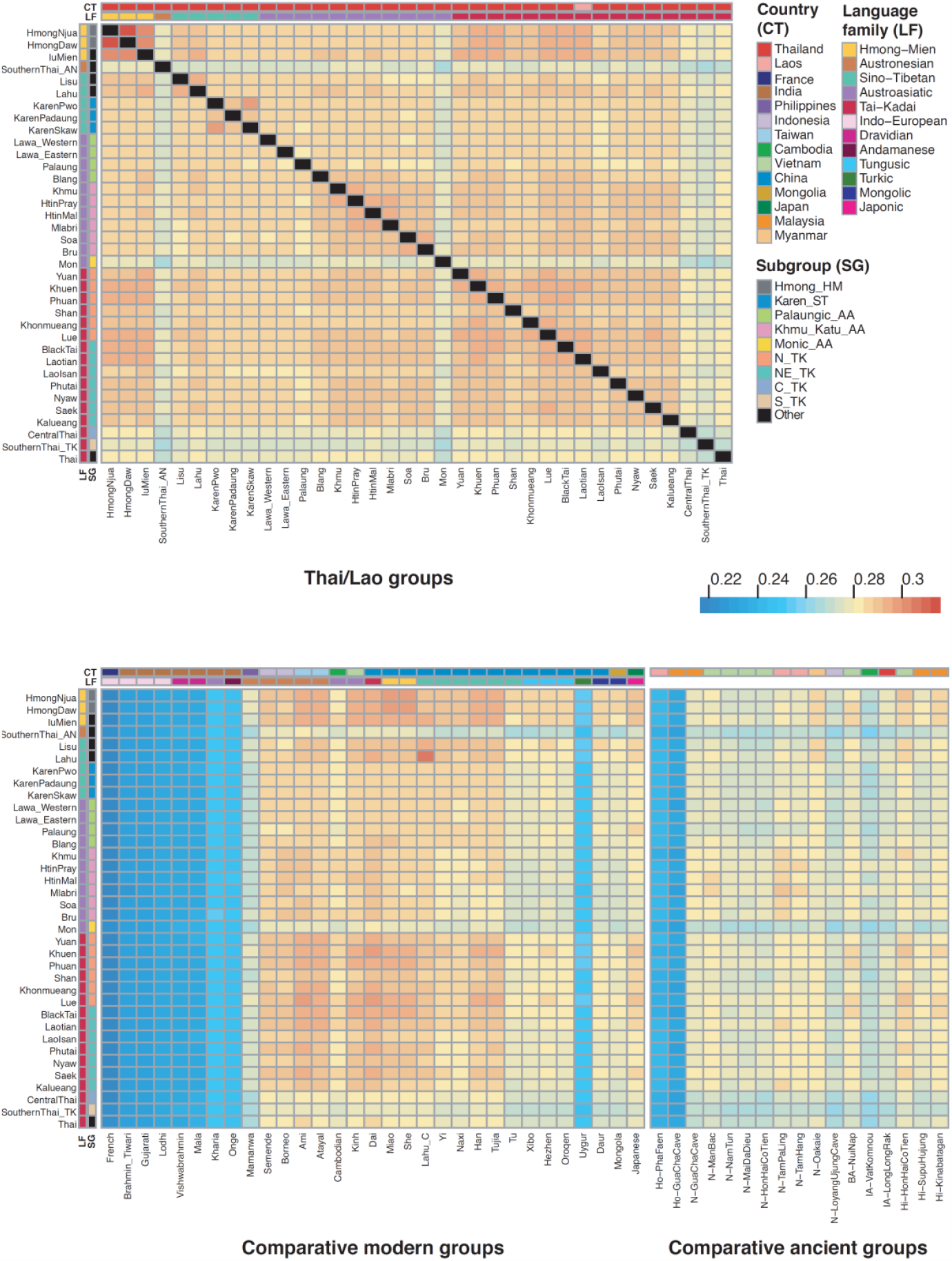
Population allele sharing profiles based on *f3* statistics. Heatmap of outgroup *f3* statistics (Thai/Lao groups, X; Mbuti) among Thai/Lao groups (upper) panel, and between Thai/Lao and other comparative modern Asian populations and ancient samples (lower). Black blocks denote missing values. The two colored bars at the top of the plot indicate the country (top) and language family (bottom) for each comparative population; and those on the side indicate language family (left) and subgroup (right) for each Thai/Lao group, according to the key at the right.

There is higher sharing between the Thai/Lao groups and other SEA and southern Chinese groups (i.e. TK, HM, and non-NEA ST Chinese groups) than with SA and NEA groups (Figure 3). The highest sharing was between Thai Lahu and Chinese Lahu. The Ami and Atayal share more alleles with the TK groups than with the SouthernThai_AN group from Thailand (Figure 3), in agreement with ADMIXTURE results (Figure 2C; Supplementary Figure 5). The ancient samples N-TamPaLing and N-TamHang share more with the Khmu_Katu_AA and NE_TK groups, but N-Oakaie shares more with the ST-speaking Lisu and Lahu groups and HM-speaking Hmong and IuMien groups. The Iron Age samples show overall less allele-sharing with Thai/Lao groups, whereas the Bronze Age and historical samples from Vietnam and Malaysia show higher sharing with the Thai/Lao TK and HM groups (Figure 3), in agreement with the ADMIXTURE results (Figure 2C).

#### ChromoPainter

To further investigate the ancestry profiles and recent past of Thai/Lao populations through haplotype-based methods, we used the ChromoPainter software (Lawson et al., 2012) and the genomes of modern Asian populations (including the Thai/Lao populations) as donors to paint the chromosomes of Thai/Lao populations. The process of “painting chromosomes” means defining the ancestry source of haplotypes along the chromosomes of a target individual by donors who share the most recent common ancestor.

We found the strongest signal is self-painting, except for the Laotian, SouthernThai_AN, SouthernThai_TK and Thai-HO which have a wider sharing profile (Figure 4A). Some finer structure within the AA groups is revealed: the Mon_AA group shows excess sharing with Indian donors; Khmu_Katu_AA groups show strong intra-group sharing but less sharing with other groups except for between the Soa and most NE_TK groups; Palaungic_AA groups show various sharing patterns, e.g. a broad sharing profile of the Blang with several other groups vs. strong self-painting only of the Palaung, and strong sharing among the Lawa_Eastern, Lawa_Western, Karen_ST groups and Shan. The relationships among Lawa, Karen_ST and Shan are also seen in PCA (Figure 2B) and ADMIXTURE results (Supplementary Figure 5). Likewise, some finer structure within the Thai TK groups is revealed: N_TK populations show strong sharing with each other and the Dai, though the Shan show additional sharing with the Lawa_Eastern and Karen_ST groups. The NE_TK groups show strong sharing with the Khmu_Katu_AA group, Cambodian, Borneo and Dai. Notably, the Laotian show a relatively broader sharing profile and high sharing with the HM groups, whereas the BlackTai show a strong self-painting profile. In addition to strong sharing with Khmu_Katu_AA groups, the C_TK group shows an excess sharing with the Indian donors, which is similar to the profile of Thai-HO. The S_TK group also shows a similar profile as C_TK but additional sharing with the AN-speaking Mamanwa, Borneo and Semende, which is similar to the profile of the SouthernThai_AN (who show even stronger and broader sharing with the other AN groups). The Thai HM groups show strong sharing with each other and the Chinese HM groups, especially the Miao. The IuMien show additional affinity to ST (especially Lahu) and N_TK groups. For the Thai ST groups, the Lisu and Lahu show strong sharing with each other and the ST-speaking Chinese Lahu, Yi and Naxi. In contrast, the Karen_ST groups show strong sharing with each other and the Lawa_Eastern, Lawa_Western and Shan.

**Figure 4.**
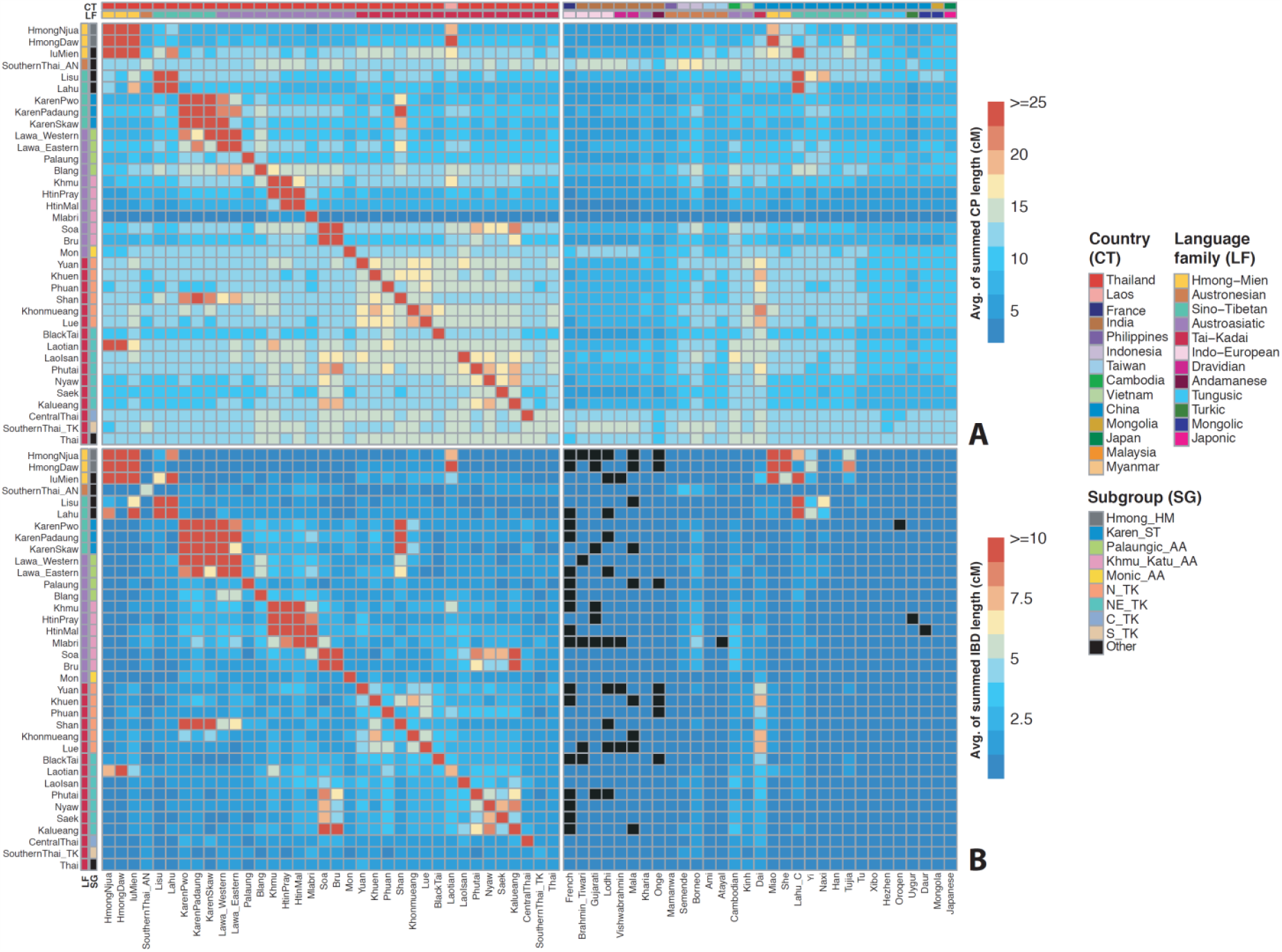
Haplotype sharing profiles as inferred by the ChromoPainter and IBD analyses. The color bars at the top denote the countries and language families while the color bars at the left denote countries and subgroups, according to the keys. (A) Heatmap of ChromoPainter results in which the recipient Y (Thai/Lao groups) is painted by donor X (Thai/Lao and other modern Asian populations), with Y denoted by each row and X denoted by each column. The heatmap is scaled by the average length in centimorgans of the summed painted chromosomal chunks of the recipient individuals from the donor individuals. (B) Heatmap of IBD sharing among Thai/Lao comparisons and between Thai/Lao and other modern Asian populations. The heatmap is scaled by the average length in centimorgans of summed IBD blocks shared between individuals from the two groups. Black blocks denote missing values.

To avoid the effects of self-painting, which is enhanced in isolated populations subject to drift, we conducted another ChromoPainter analysis in which we excluded individuals sampled from this study as donors. The three Thai groups from previous studies, HtinMal, Mlabri, and Thai-Ho, were still included as donors but were removed from being recipients, in order to capture some local ancestry from Thailand (Supplementary Figure 6). With self-painting not allowed, sharing profiles with the comparative Asian populations become more pronounced. In particular: the profile of the Palaung becomes more similar to other Palaungic_AA groups; all of the Khmu_Katu_AA groups are highly painted by the HtinMal and Mlabri donors (who are also Khmu_Katu_AA groups), suggesting strong affinities among the Khmu_Katu_AA groups; the sharing profile of BlackTai with other groups is revealed to be similar to both N_TK (sharing with the Dai) and NE_TK (sharing with Borneo) groups. Previously-identified sharing profiles also become more obvious, e.g. high sharing between N_TK and Dai donors, NE_TK and AA donors (e.g. Cambodian and Kharia), and C_TK and S_TK and Indian donors.

#### Identity by descent (IBD)

The IBD analysis generally captured the main features of the ChromoPainter results with less resolution for the sharing with populations outside Thailand/Laos (Figure 4B). However, the length of shared IBD segments provides a rough time frame for the interactions within/between populations (Ralph and Coop, 2013; Al-Asadi et al., 2019), and the number and length of IBD segments shared within a population can be used to infer population demography (Browning and Browning, 2015; Browning et al., 2018; Ceballos et al., 2018; Severson et al., 2019). We found that all AA (except for Mon and Blang), HM (except for IuMien), and ST groups exhibit high within-population IBD sharing (Supplementary Figure 7), with the Mlabri showing the greatest levels by far of within-group IBD sharing, in agreement with their enhanced self-painting in the ChromoPainter analysis (Figure 4A). Most of these groups are hill tribes, suggesting strong drift effects in isolated groups in this remote mountainous area. Low levels of within-group IBD sharing, suggesting either population expansion or admixture, is observed in most TK and AN groups, who mostly occupy the lowlands and tend to exhibit broader sharing profiles in the ChromoPainter analyses (Figure 4A; Supplementary Figures 6-7).

The IBD sharing between populations was broken down into categories based on the length of shared IBD blocks, in order to infer the approximate time of interactions; the longer the shared IBD blocks, the more recent the interaction as there has been less time for recombination to shorten the IBD blocks. We analyzed three categories of IBD blocks: 1-5 cM, 5-10 cM, and >10 cM (Supplementary Figure 8); these correspond very roughly to time intervals of 1,500-2,500 ya, 500-1,500 ya, and 0-500 ya, respectively (Ralph and Coop, 2013). Overall, all populations show some sharing with other populations, and most of the Thai/Lao groups share IBD blocks during the 1,500-2,500 ya interval. In general, shared IBD was restricted to populations from the same language family, as reflected in Figure 4B: the Thai/Lao TK and AA populations share IBD segments with TK-speaking Chinese Dai and AA-speaking Cambodian, respectively; Thai HM populations share IBD segments with the HM-speaking Miao and She from China; and Thai ST groups share IBD segments with ST-speaking groups from China. Interestingly, an exception to this pattern of shared IBD restricted to populations from the same language family occurs in southern Thailand, where both SouthernThai_TK and SouthernThai_AN groups share IBD segments with the AA-speaking Mlabri, although the SouthernThai_AN additionally share IBD segments with AN-speaking groups from Sumatra and Borneo. The pattern becomes much more localized in later periods, with sharing restricted to a few groups in northern and northeastern Thailand (Supplementary Figure 8).

We also estimated recent changes in effective population size within the past 50 generations using the IBD sharing within each individual population (Browning and Browning, 2015) (Supplementary Figure 9). Most populations show a decline around 20 generations ago that is followed either by a constant population size or a small increase, but SouthernThai_TK, SouthernThai_AN, and Thai-HO show population increases only beginning around 10-20 generations ago. This result emphasizes the difference between populations from southern/central Thailand vs. those from northern/northeastern Thailand and Laos. However, we caution that our estimation of effective population size is likely to be uncertain for populations with large effective population sizes in recent generations, due to the assumption of a constant growth rate and insufficient sample sizes for accurate estimation (Browning and Browning, 2015; Browning et al., 2018).

#### Investigating shared ancestry with f4-statistics

The *f4*-statistics of the form *f4* (W, X; Y, Outgroup) were used to formally test whether population W or X shares more ancestry with population Y. We first investigated the relationships among Thai/Lao groups from the same language family/subgroup by computing *f4*-statistics of the form (group 1, group 2; group 3, Mbuti), where group 1 and group 2 are from the same language family/subgroup while group 3 is from a different language family/subgroup. By convention, a Z-score > 3 or < −3 indicates that group 3 shares significant excess ancestry with group 1 or 2, respectively; nonsignificant Z-scores indicate that group 1 and 2 form a clade and share equivalent amounts of ancestry with group 3. The results indicate that there is no significant sharing of ancestry between HM or ST groups (except for Lahu, and Lisu with HM groups) and non-HM or non-ST groups, respectively (Supplementary Figure 10A-B; Supplementary Table 2). However, there are numerous instances of an AA or TK group sharing excess ancestry with a non-AA or non-TK group (Supplementary Figure 10C-10E; Supplementary Table 2); this heterogeneous ancestry sharing profile also reflects the putative South Asian ancestry in some AA and TK groups (Supplementary Figure 10C and 10E; Supplementary Table 2). In particular, the profiles of NE_TK and N_TK groups show strong excess sharing with each other and the HM groups, followed by ST and AA groups (Supplementary Figure 11A-11C; Supplementary Table 3). Many of the highest Z-scores come from comparisons involving the Laotian population (Supplementary Figure 10D and 11A; Supplementary Tables 2-3), in agreement with their broader haplotype sharing profiles (Figure 4). The profiles of Khmu_Katu_AA and Palaungic_AA exhibit excess sharing with each other and higher excess sharing with the Karen_ST groups than with the other ST groups, which is also consistent with the haplotype sharing profiles (Figure 4; Supplementary Figure 11D-11E; Supplementary Table 3). In addition, we found that Thai-HO and CentralThai form a clade in all the tests (Z scores within +/- 1.5), suggesting their close relationship in agreement with previous analyses (Supplementary Figure 10E; Supplementary Table 2).

We further investigated whether any of the Thai/Lao groups share excess ancestry with representative East Asian groups, compared to Han Chinese, by computing *f4*-statistics of the form (East Asian group, Han Chinese; Thai/Lao group, Mbuti). A Z-score > 3 indicates that the Thai/Lao group shares excess ancestry with the East Asian groups, while a Z-score < −3 indicates that the Thai/Lao group shares excess ancestry with Han Chinese; nonsignificant Z-scores indicate no excess ancestry sharing of the Thai/Lao group with either the East Asian group or Han Chinese. Based on the allele and haplotype sharing profiles (Figures 3-4), we used Atayal, Dai, Cambodian, Miao and Naxi as representative groups speaking AN, TK, AA, HM and ST languages, respectively. Almost all of the Thai/Lao TK groups and the SouthernThai_AN population share excess ancestry with Atayal and Dai (Supplementary Figure 12), share more ancestry with Han than with Cambodian or Naxi (although the SouthernThai_AN shares less excess ancestry with Cambodia than other Thai/Lao groups), and show either a slight excess sharing, or no excess sharing, with Miao. These results provide further support for a genetic relationship between TK and AN groups. In addition, the grouping among AA Thai/Lao groups was also supported by this test; the Monic_AA show excess sharing only with the Dai, while the Khmu_Katu_AA and Palaungic_AA groups are distinguished by the former sharing excess ancestry with Atayal and having no significant Z-scores with Cambodian vs. Han, while the latter have no significant Z-scores with Atayal and share excess ancestry with Han when compared with Cambodian. These results suggest more AN/TK and AA related ancestry in the Khmu_Katu_AA group, and more Han related ancestry in the Palaungic_AA group. The ST and HM populations are similar in their overall patterns to the Palaungic_AA group, except that the HM populations share the most excess ancestry with the HM-speaking Miao, while the ST populations share less excess ancestry with Han than do most of the other Thai/Lao groups when compared to the ST-speaking Naxi.

We next used *f4* (Thai/Lao group, Han; Indian group, Mbuti) to investigate the putative South Asian-related admixture shown by PCA and ADMIXTURE results (Figure 2), and the haplotype sharing profiles (Figures 4A). Several TK and AA Thai/Lao groups share significant excess ancestry with the AA-speaking Kharia (Supplementary Figure 13). By contrast, the Mon, SouthernThai_TK and SouthernThai_AN share excess ancestry with every other Indian group (but not the Kharia or Onge), and they are the only Thai/Lao groups to share excess ancestry with the other Indian groups. They are also the only groups (along with CentralThai) that share less ancestry with Onge than do Han. These results highlight the distinctive nature of the Indian-associated ancestry in the Mon and southern Thai groups, compared to other Thai/Lao groups.

We also performed an *f4* analysis of the form *f4* (ancient samples, Han; Thai/Lao groups, French), with only transversions (3,090-53,870 SNPs), to assess allele-sharing between the Thai/Lao groups and the ancient samples (Supplementary Figure 14). Most populations show no significant differences in ancestry sharing with the Hòabìnhian samples vs. Han Chinese, except that the Mon and SouthernThai_TK share more alleles with Han while Blang shares more allele with Ho-PhaFaen. Many of the Thai/Laos populations show significant ancestry sharing with most of the Neolithic samples; however, the Mon_AA, C_TK, S_TK, and SouthernThai_AN groups share excess ancestry with Han compared to the ancient samples, and this pattern becomes weaker in later periods.

### Population histories investigated by admixture graphs

Constructing admixture graphs, using either a combination of *F*-statistics or a covariance matrix of the allele frequencies, is another method to explore the shared genetic ancestry, admixture events and historical population divergence among multiple populations simultaneously (Nielsen, 2018). TreeMix (Pickrell and Pritchard, 2012) and AdmixtureBayes (Nielsen, 2018) analyses were first carried out to survey the potential admixture graphs based on the covariance matrix of allele frequencies, and then qpGraph (Patterson et al., 2012) was used to further test if these graphs provide a reasonable fit to the data, using a combination of *F*-statistics.

We began with a maximum-likelihood tree inferred by TreeMix with Mbuti (as the outgroup), French, South Asians (N_Indian and Onge), representative East Asian groups (same as those used in the *f4* analyses), ancient samples with more than 130,000 overlapping SNPs (<65% missing data; these are Ho-PhaFaen, N-TamPaLing, N-GuaChaCave, IA-LongLongRak, and Hi-Kinabatagan), and Thai/Lao groups. The N_Indian, TK, AA, Hmong_HM, and Karen_ST groups were grouped based on linguistic classification and ChromoPainter results (see Materials and Methods). The overall topologies with and without migration are similar, except for shifts involving a few groups (Supplementary Figure 15A). The SouthernThai_AN, S_TK, Monic_AA, C_TK and Thai-HO, together with the ancient samples, fall outside a clade containing the remaining Thai/Lao groups and the representative East Asian groups.

The standard error of the residuals decreases from 15.6 to 12.3 when adding 3 migration events (Supplementary Figures 15B) and all groups from the same language family now form a clade except that the Karen_ST is placed in the AA clade together with Neolithic/Iron Age samples (N-GuaChaCave, N-TamPaLing, and IA-LongLongRak); the AN-speaking Atayal falls in the TK clade; and the Southern Thai_AN is placed in between the Hòabìnhian-related Onge/Ho-PhaFaen and the historical Hi-Kinabatagan samples. There were three migrations inferred: one from N_Indian to Mon_AA and IA-LongLongRak; one from the ancestor of all samples after the divergence of N_Indian and French to S_TK, C_TK, and Thai-HO; and one from the Hòabìnhian sample to the Neolithic samples.

To investigate the genetic ancestry in each language family, we built admixture graphs using AdmixtureBayes, and then further investigated these admixture graphs with qpGraph (Figure 5). To begin with, we built a backbone admixture graph with the outgroup Mbuti, N_Indian, and the representative East Asian groups (Figure 5A); the first split separates the N_Indian from the East Asian groups, then the Naxi are separated from the other groups. The ancestor of Atayal and Dai is admixed from ancestors of N_Indian and Miao with 6% and 94% ancestry, respectively. The ancestor of Cambodian is admixed with 73% ancestry from the ancestor of Dai and 27% from the ancestor of all East Asian groups. We then explored graphs for groups from each language family. For the SouthernThai_AN group (Figure 5B), the Indian-related ancestor contributes 27% ancestry to the SouthernThai_AN, with the remaining 73% contributed by an admixed ancestor with AA- and AN-related ancestry. For the four TK groups (Figure 5C), the NE_TK and N_TK groups are in the same clade, and this clade contributes 88% to C_TK and 83% to S_TK. The remaining ancestry for C_TK and S_TK is contributed by Indian-related ancestry, which reflects SA-related admixture that is consistent with previous results (Figures 2 and 4A; Supplementary Figure 13). This graph does not include any EA source populations as their inclusion leads to unacceptable graphs (worst-fitting Z = −7.037; Supplementary Figure 16), probably because the Dai have broad attraction to all the TK groups as well as Atayal and Cambodian, as most of the outlier Z-scores involve the Dai. However, this graph still provides essentially the same topology for the TK groups as in Figure 5C with the N_TK now forming a clade with the Dai and Atayal while the NE_TK share more ancestry with Cambodian. To reduce complexity/redundancy in the modelling, we did not include the Thai-HO in the graph as their ethnolinguistic background is unclear and their genetic profile is very similar to C_TK (Supplementary Figure 10E; Supplementary Table 2). The graph of AA groups (Figure 5D) includes several admixture events, and indicates that the Khmu_Katu_AA and Palaungic_AA subgroups are more closely-related, while the Monic_AA subgroup is distinguished from these by N-Indian-related ancestry, in agreement with previous results (Figures 2 and 4A; Supplementary Figure 13). For the HM groups (Figure 5E), there is a divergence between the Dai and a Miao-Hmong clade, while the IuMien are admixed with 29% ancestry from an ancestor of the Hmong and 71% from an ancestor of the Dai. The additional TK-related ancestry in IuMien is consistent with haplotype-sharing and *f4* results (Figure 4; Supplementary Figure 12). The graph of ST groups indicates that Lisu, Lahu and Naxi form a clade, while the Karen_ST have additional Cambodian-related ancestry (Figure 5F); this AA-related admixture in the Karen is in agreement with the haplotype-sharing and Treemix results (Figure 4, Supplementary Figure 15).

**Figure 5.**
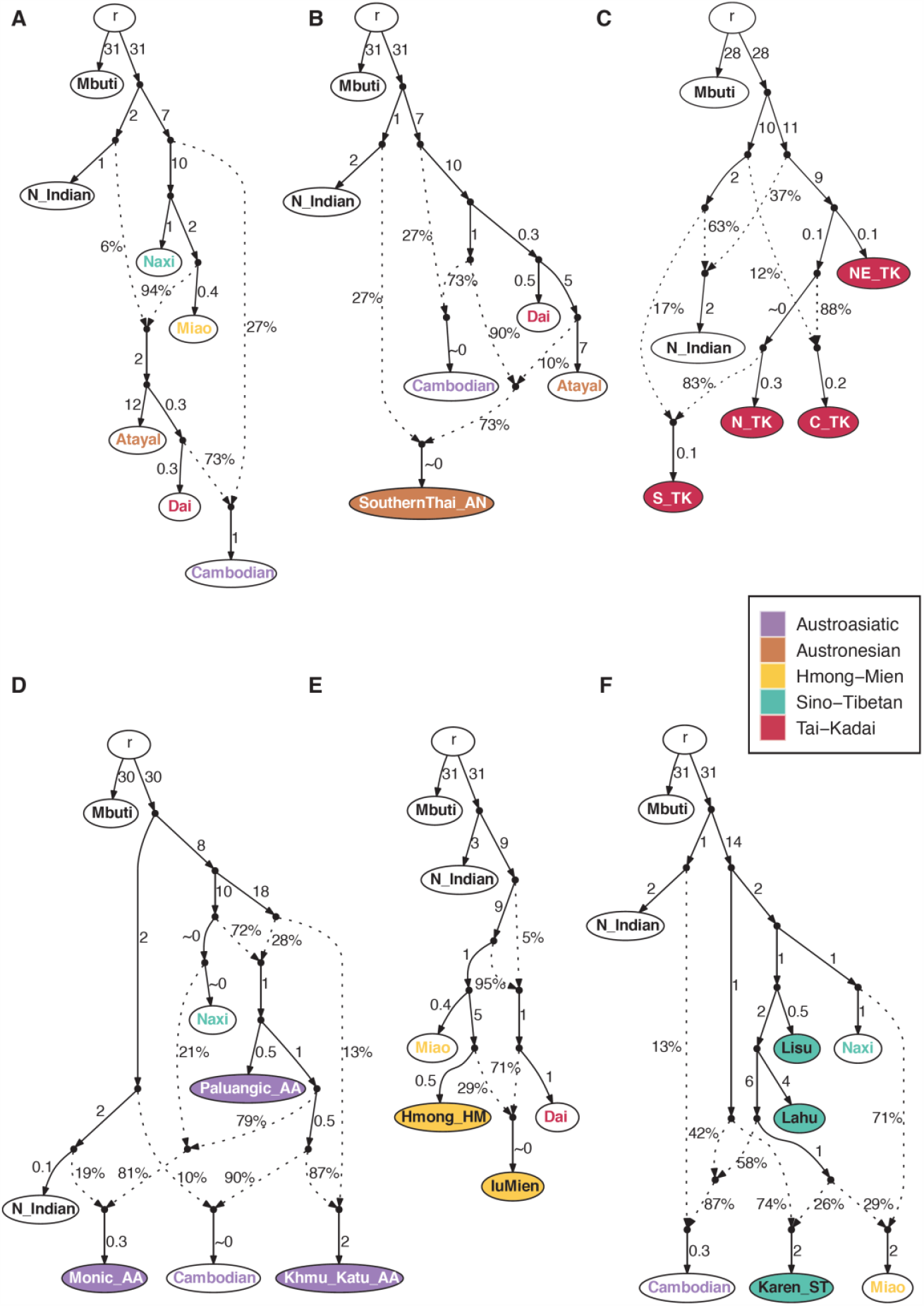
Admixture graphs for the Thai/Lao groups, for each language family. The node r denotes the root. White nodes denote backbone populations. Backbone population labels and Thai/Lao nodes are colored according to language family. Dashed arrows represent admixture edges, while solid arrows are drift edges reported in units of FST×1,000. (A) backbone populations (worst-fitting Z = 0.861). (B) AN group (worst-fitting Z = −1.713). (C) TK groups (worst-fitting Z = −2.270). (D) AA groups (worst-fitting Z = 2.101). (E) HM groups (worst-fitting Z = −2.028). (F) ST groups (worst-fitting Z = −2.873).

### South Asian-related admixture investigation

The results of PCA, ADMIXTURE, ChromoPainter, *f4*-statistics, and admixture graph analyses (Figures 2, 4-5; Supplementary Figures 13 and 15) all suggest South Asian related ancestry in the Mon, SouthernThai_AN, SouthernThai_TK, CentralThai, and Thai-HO. To further analyse the details of this putative admixture, we used the GLOBETROTTER software (Hellenthal et al., 2014), based on the output of ChromoPainter, to infer the number of admixture events, identify proxies for the admixture sources, and date admixture events. Again, to reduce redundancy in the modelling, we did not include the Thai-HO in the graph as their ethnolinguistic background is unclear and their genetic profile is very similar to C_TK (Supplementary Figure 10E; Supplementary Table 3). We included Yuan in the source estimation as a control because they did not show any SA-related admixture signal but are geographically close to the other groups. For each group (including the Yuan control group), a single admixture event is inferred (Figure 6A). However, the admixture inferred for the Yuan is statistically uncertain, and the composition of sources is quite different compared to the sources inferred for the other groups: the dominant major sources are 46% from AA-speaking Kinh and 35% from TK-speaking Dai while the dominant minor sources are 4% from Indian Gujarati and 2% from ST-speaking Naxi. For the other groups, the dominant proxy for the major source is the Kinh, ranging from 45% to 63% (and 7-11% for the Dai), with the minor source from the Indian Brahmin Tiwari (10%) for the SouthernThai_TK and Gujarati (7-18%) for the rest. Apart from the dominant sources, the SouthernThai_AN are also inferred to have more AN-related (Mamanwa, Borneo, Semende, Atayal, and Ami) ancestry (19% vs. 9% in SouthernThai_TK and below 5% in the others), while the Mon have more ST-related (Lahu, Naxi, and Yi) ancestry (9% vs. below 4% in the others), in agreement with the admixture graphs (Figure 5).

**Figure 6.**
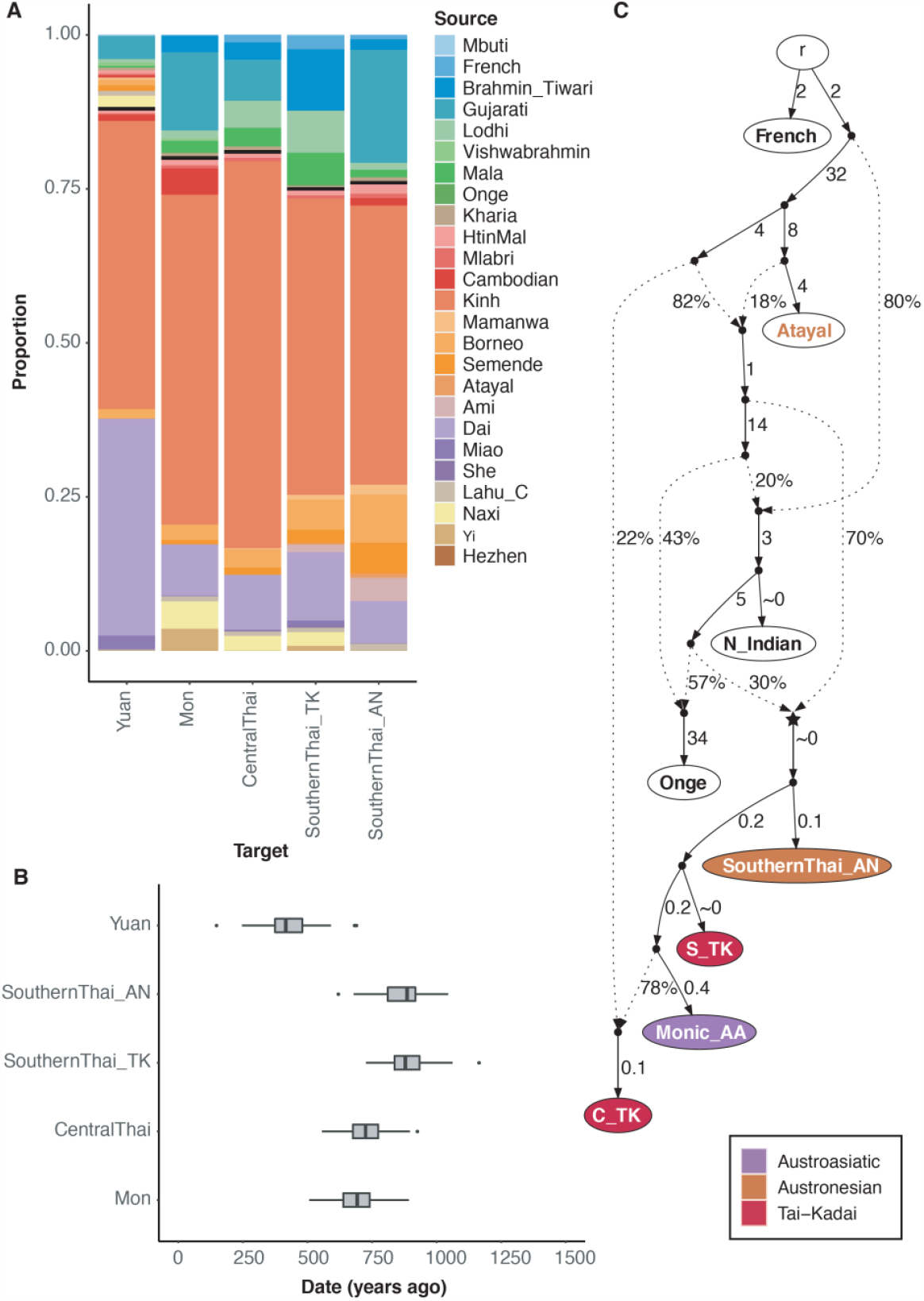
Investigation of putative SA-related admixture. (A) GLOBETROTTER estimation of admixture sources for four Thai groups (Mon, Central Thai, SouthernThai_TK and SouthernThai_AN) with putative SA-related ancestry, and for the Yuan group as a control without putative SA-related ancestry. Different sources are denoted by different colors. (B) GLOBETROTTER estimates of the admixture date in the SA-influenced Thai groups. Results are based on 100 bootstraps. (C) Admixture graph for the Thai groups with SA-related admixture (worst-fitting Z = −1.646). The node r denotes the root. White nodes denote backbone populations. The star-shaped node denotes the N_Indian-related source contributing to all of the SA-related Thai groups. Backbone population labels and Thai nodes are colored according to language family. Dashed arrows represent admixture edges, while solid arrows are drift edges reported in units of FST×1,000.

We next estimated the admixture dates using GLOBETROTTER; these range between 600-900 ya for the SA-related populations with the dates for both southern Thai populations tending to be older than those for the other groups (Figure 6B). We also estimated the admixture date for the Yuan even though the admixture is uncertain; a much younger date was inferred (∼400 ya). We also used another admixture dating software, ALDER, that is based on the decay of linkage disequilibrium (LD) (Supplementary Figure 17), which gave results overall falling in the similar time range with a slightly younger distribution of dates (500-750 ya). We used the most dominant major (Kinh) and minor (Gujarati) sources inferred by GLOBETROTTER as sources for ALDER. However, the LD decay curves of all the groups could not be fitted with the Kinh LD curve, while the Gujarati LD curve provided a fit for the SA-related groups but not for the Yuan. The ALDER dating was therefore carried out using just the Gujarati LD curve.

Finally, we also built an admixture graph for the Thai groups with inferred SA-related ancestry (Figure 6C). We included for comparison French (as the outgroup), N_Indian, and Onge to investigate if the SA-related source is most similar to European, northern Indian, or southern Indian ancestries, and we also included Atayal as a source of East Asian ancestry. An acceptable graph (worst-fitting Z = −1.646) indicates that the SA-related ancestry traces back to a single ancestral node (the star node in Figure 6C) that contributes 30% to the ancestry of the SA-related Thai groups, which is similar to the amount of SA-related source (minor source) estimated from GLOBETROTTER (Figure 6A). The C_TK are inferred to have an additional 22% ancestry from a lineage related to Atayal, similar to other admixture graphs for TK groups (Figure 5A, Supplementary Figure 16). Inclusion of more EA source populations and using Mbuti as an outgroup does not provide an acceptable graph (worst-fitting Z = −4.110; Supplementary Figure 18) but the overall topology is consistent with that in Figure 6C. While an AA-related ancestor contributes more than 80% ancestry to the SA-related Thai groups, suggesting that they are all mainly AA-related despite some of them speaking TK or AN languages, additional ancestry comes from TK, N_Indian, and Onge sources.

## Discussion

Previous detailed genetic studies of Thai/Lao populations focused primarily on uni-parentally inherited markers and found: contrasting patterns of paternal vs. maternal genetic variation in hill tribe and hunter-gatherer groups (Oota et al., 2001; Besaggio et al., 2007; Kutanan et al., 2018a and b; Kutanan et al., 2019); more ancient lineages and heterogeneity of the AA-speaking groups (Kutanan et al., 2017); genetic relatedness between central Thais and AA-speaking Mon with both showing South Asian specific haplogroups (Kutanan et al., 2018b; Kutanan et al., 2019); and relatedness between TK and AN speaking groups (Kutanan et al., 2018b) that is also supported by a recent ancient DNA study (Yang et al., 2020). However, additional insights into the genetic history of this region, e.g. fine-scale structure, the extent and dating of South Asian admixture, and other population interactions have not been investigated. Here, we analyzed genome-wide SNP data from 36 populations encompassing all five major linguistic families from Thailand and Laos. Our major findings, which we discuss below, are: genetic clustering and heterogeneity of AA speaking groups; the genetic structure of the hill tribes; differences among the four major TK speaking groups according to geographic region; and South Asian admixture.

### Genetic heterogeneity of Austroasiatic speaking populations in Thailand

AA speakers (comprising ∼102 million people speaking 167 languages) are widespread across Asia, from South Asia (Bangladesh and India) to southern China and MSEA (Eberhard, Simons and Fennig 2020). Although there were two competing hypotheses of AA origins that are related to rice cultivation, i.e. South vs. Southeast Asian origins (Chaubey et al., 2011; Diffloth 2005), the latter is supported by genetic evidence (Chaubey et al., 2011). The AA people in SEA are most likely related to farmers who knew rice and millet cultivation and moved from their homeland, probably located near the Yangtze River, to the coast and then down the rivers of mainland China to SEA ∼4 kya (Weber et al., 2010; van Driem, 2017; Lipson et al., 2018; McColl et al., 2018). However, prior to the movement of prehistoric AA-related groups southward, present-day MSEA (both upland and lowland) was home to hunter-gatherers whose descendants are genetically related to groups in southern Thailand and west Malaysia, such as the Maniq and Jehai (Jinam et al., 2012). The Neolithic farmer expansion did not completely replace the hunter-gatherers but admixed with some of them, as reflected by both ancient and modern DNA studies (Lipson et al., 2018; McColl et al., 2018; Kutanan et al., 2017; Liu et al., 2020).

Previous genetic and linguistic evidence suggested heterogeneity of the Thai AA people (Xu et al., 2010; Kampuansai et al., 2017; Kutanan et al., 2017; Eberhard, Simons and Fennig, 2020) but further genetic groupings have not yet been investigated. In this study, several lines of evidence indicate that the Thai AA speaking populations fall into 3 primary groups: Monic_AA, Khmu_Katu_AA and Palaungic_AA (Figures 2-4; Supplementary Figure 12). The language of Mon is in the Monic branch, the sister clade of Aslian and Nicobarese, while the linguistic branch of Khmu_Katu_AA groups are Khmuic for HtinMal, HtinPray, Mlabri and Khmu, and Katuic for Soa and Bru; the Palaungic branch includes languages of the Lawa_Eastern, Lawa_Western, Palaung and Blang. In contrast to linguistic studies placing Khmuic and Palaungic languages in the same clade (Diffloth, 2005), we find a closer relationship between populations who speak Khmuic and Katuic, which might be explained by the concept of center of gravity (Blench, 2015). This idea proposes that after the Neolithic expansion of AA ancestors from southern China to MSEA, early AA speakers were concentrated along the middle Mekong in present-day northern Laos. Some groups subsequently moved westward and were the ancestors of Palaungic and Monic groups, and during this process they came into contact with other different linguistic groups (e.g. Mon with Burmese ancestors, Lawa_Eastern and Lawa_Western with Karen_ST, and Palaung with ST groups from NEA), as shown by population structure and relationship analyses and *f4* tests (Figures 2-4; Supplementary Figure 11; Supplementary Table 3). These different contact histories would promote subsequent differentiation of the Palaungic and Monic groups from their Khmuic and Katuic ancestors. Meanwhile, the Khmuic and Katuic ancestors might have moved up and down the Mekong and had more contact with each other, thus accounting for their closer genetic relationship with each other. In this region, the Khmuic and Katuic speaking people may have also interacted with TK groups in Laos and Northeastern Thailand and promoted their genetic affinity (Figures 2B, 3-4; Supplementary Table 3). However, some differentiation between the Khmuic and Katuic groups can be seen in the haplotype sharing (Figure 4) and ADMIXTURE results for *K*=10 (Supplementary Figure 5). Additional studies of AA groups from Thailand, e.g. Pearic and Khmer speaking groups and other MSEA countries are needed to provide more insights into the genetic structure of AA-speaking people.

### The hill tribes

Consisting of ∼700,000 people, there are nine officially recognized hill tribes in Thailand: the AA-speaking Lawa (Lawa_Eastern and Lawa_Western), Htin (HtinMal and HtinPray) and Khmu; the HM-speaking Hmong (HmongNjua and HmongDaw) and IuMien; and the ST-speaking Karen (KarenPwo, KarenPadaung, and KarenSkaw), Lahu, Lisu and Akha (Schliesinger, 2000, 2001; Penth and Forbes, 2004). Living in a remote and isolated region of Thailand, the hill tribes are of interest for their cultural variation in residence pattern after marriage, i.e. patrilocality vs. matrilocality (Oota et al., 2001; Besaggio et al., 2007; Kutanan et al., 2019, 2020).

Most of the hill tribes are isolated from the lowlanders and from each other, which enhances genetic drift and inbreeding, as found in previous studies of autosomal STR (Kampuansai et al., 2017) and mtDNA and MSY variation (Kutanan et al., 2020). We therefore expected similar indications of isolation in our study, which included eight of the official hill tribes (all but the Akha). Indeed, we found four groups with their own ancestry components in the ADMIXTURE results at *K* = 10 (Supplementary Figure 5): Lahu (light green), Karen_ST (grey), Htin (Mal and Pray) and Khmu (mint) and Hmong_HM (peach), in agreement with their higher IBD sharing within groups (Supplementary Figure 7). In contrast, the Lawa (Eastern and Western), Lisu and IuMien do not stand out in the ADMIXTURE analysis, and they have relatively less within group IBD sharing (Supplementary Figure 7), show excess allelic sharing with many other populations in the *f4* results (Supplementary Tables 2-3), and shared haplotypes with other groups (Figure 4A; Supplementary Figure 6). These results indicate that not all hill tribes can be characterized simply by high degrees of isolation and genetic drift; the Lawa, Lisu, and IuMien instead seem to have had more interactions with other groups, and so we will focus further discussion on these three hill tribes. The Lawa (Eastern and Western) are the native groups of northern Thailand and inhabited lowland areas before some of them moved to the highlands (Lawa_Western) while others remained in the lowlands or mid-lands (Lawa_Eastern) (Nahhas, 2007). By contrast, the Karen in Thailand are refugees who claim to be the first settlers in Myanmar before the arrival of Mon and Burmese people, and moved from Myanmar beginning around 1750 A.D. due to the growing influence of the Burmese (Kuroiwa and Verkuyten, 2008; Gravers, 2012). The Lawa share ancestry with the Karen_ST (Figure 4; Supplementary Figure 5), in agreement with previous findings of shared MSY haplotypes (Kutanan et al., 2020). Genetic relatedness between Karen and Lawa groups was also reported in a previous genome wide study (Xu et al., 2010). In northern Thailand, Lawa and Karen had been historically contacted since ∼ the 13^th^ century A.D. during the Lanna Period (Lewis and Lewis, 1984). Because the languages of AA-speaking Lawa and ST-speaking Karen are different, geographic proximity along the border between northern/northwestern Thailand and Myanmar is the most likely factor that promoted admixture between these groups.

The Lisu and the Lahu are originally from southern China, and speak closely related languages that belong to the Loloish branch of ST (Bradley, 1997). Shared genetic ancestry between Lisu and Lahu is evident in the haplotype sharing and admixture graph results (Figure 4 and 5F; Supplementary Figure 15), although there are differences: Lisu have mixed ancestries probably due to Sinicization in southern China before movement to Thailand (Schliesinger, 2000) or interactions with northern Thai lowlanders after settlement in Thailand (Penth and Forbes, 2004), while the Lahu are more isolated, e.g. the ADMIXTURE result for *K* = 7 (Supplementary Figure 5) and the IBD sharing results (Supplementary Figure 7), in agreement with a previous study of uniparental markers (Kutanan et al., 2020). There is strong ancestry sharing between the Thai Lahu and Chinese Lahu (Figures 3-4), and the Chinese Lahu are moreover genetically similar to Vietnamese Lahu (Liu et al., 2020), indicating a close relationship among Lahu from MSEA and China.

Although the IuMien and Hmong are descended from proto-HM groups from central and southern China (Wen et al., 2005) and are linguistically related, they behave differently in many analyses (Figures 3-5 Supplementary Figures 6 and 12). The Hmong show genetic signatures of isolation, such as higher IBD sharing within groups (Supplementary Figure 7), in agreement with a previous study of uniparental markers (Kutanan et al., 2020), whereas the IuMien show affinities not only with the Hmong, but also with TK speaking groups and ST speaking Lahu from both Thailand and China (Figure 4). The differential affinities of HM groups to TK and ST groups has also been shown in two recent genome-wide studies (Liu et al, 2020; Xia et al., 2019). In addition, the sharing of features between IuMien (but not Hmong_HM) and Sinitic languages (Blench, 2008) indicates that IuMien similarities with other East Asian populations is evident both genetically and linguistically. The higher genetic isolation of the Hmong could reflect cultural isolation arising from a strong preference for marriage within Hmong groups, while the lower genetic isolation of the IuMien could reflect the pronounced IuMien cultural preference for adoption (Schliesinger, 2000; Jonsson, 2005; Besaggio et al., 2007).

Though the Mlabri are not officially regarded as a hill tribe, this minority group lives in the mountainous area and is of interest due to their unique hunting-gathering life style, enigmatic origin, and very small census size (∼400 individuals) (Eberhard, Simons and Fennig, 2020). The Mlabri language belongs to the Khmuic branch of AA languages that is also spoken by their neighbors, Htin (Mal and Pray subgroups) and Khmu, suggesting shared common ancestry, and oral tradition indicates that the Htin are the ancestors of the Mlabri (Oota et al., 2005). A previous genome-wide study also supported genetic affinities between the Mlabri and the HtinMal (Xu et al., 2010), while uniparental studies show different affinities. One the paternal side (MSY), Mlabri HtinMal, HtinPray and Khmu show genetic relationships, consistent with the oral tradition, while on the maternal side (mtDNA) Mlabri shows genetic relationships with the Katuic-speaking Soa and Bru from northeastern Thailand (Kutanan et al., 2018a). Our present results also support genetic relatedness among Mlabri, Htin (Mal and Pray), Khmu, Soa and Bru within the Khmu_Katu_AA group (Figure 2B; Supplementary Figures 5-6). The Mlabri, Htin, Khmu, Soa and Bru all migrated from Laos about 100-200 years ago (Schliesinger, 2000), thus close relatedness among them might reflect gene flow among various groups in Laos before their independent migrations to Thailand. However, the Mlabri stand out among these groups in exhibiting extremely high levels of within-group IBD sharing (Supplementary Figure 7), indicating strong genetic drift and isolation, consistent with previous investigations of mtDNA, Y chromosome, and autosomal diversity (Oota et al., 2005; Xu et al., 2010; Kutanan et al., 2018a). Moreover, the IBDNe software failed to estimate the population size, probably also due to their extremely high within-group IBD sharing. Both the small census size and recent origin within the past 1000 years (Oota et al., 2005; Kutanan et al., 2010), combined with geographic isolation, could account for the very low genetic diversity of this group.

### Regional variation of Tai-Kadai speaking populations

With an origin from south/southeastern China (Sun et al., 2013; Pittayaporn, 2014), the TK language family comprises around 95 languages spoken by ∼80 million people in northeast India, southern China, Vietnam, Myanmar, Cambodia, Thailand and Laos (Eberhard, Simons and Fennig 2020). The TK languages spread to MSEA around 1-2 kya (Pittayaporn, 2014), and previous genetic studies estimated an expansion time for TK groups ∼2 kya (Kutanan et al., 2019) and found relatedness between modern TK populations and ancient Iron Age samples (McColl et al, 2018). MtDNA and MSY data indicate contrasting genetic variation and genetic differences between major TK groups in the North, Northeast and Central regions of Thailand (Kutanan et al., 2019), suggesting different migration routes of TK groups expanded from China. A previous genome-wide study also reported substructure of Thais in each region (Wangkumhang, 2013), however, these previous studies did not investigate this substructure in detail. In this study, although there is genetic homogeneity of TK groups compared with groups speaking other languages (i.e. AA and ST languages), and allelic sharing among N_TK and NE_TK groups (Supplementary Figures 10-11; Supplementary Tables 2-3), overall we find fine structure of TK groups in each geographic region (Figures 2B, 3-4; Supplementary Figures 5-6) that primarily reflects heterogeneity in admixture with local AA groups and geographic proximity. Northern Thailand is close to southern China; the N_TK groups are genetically close to the southern Chinese Dai and less mixed with local AA in the region. In contrast, Northeastern Thailand shares a border to Laos; the NE_TK groups are more related to the Khmu_Katu_AA groups that are widely distributed in Laos and recently migrated to Thailand. Central and southern Thailand share a border with Myanmar to the west; the central Thais (C_TK) and southern Thais (S_TK) have close genetic relationships with the Mon, who migrated from Myanmar.

Additionally, the N_TK groups are genetically closer to the clade of TK-speaking Dai and AN-speaking Atayal in the admixture graph (Supplementary Figure 16). This supports a common origin of TK and AN language families in southern China, as suggested previously based on linguistic and genetic evidence (Thurgood, 1994; Sagart, 2004; Kutanan et al., 2018b; Yang et al., 2020), as well as less contact of the N_TK groups after their split from the TK-AN source from southern China. Overall, our results indicate diversity of Thai TK populations, and so future whole genome or genome-wide studies should include a geographically-representative sample of Thai TK groups, to fully capture this diversity. In addition, our results provide insights into the relationships of the Thai-HO group, which was published earlier but without any details concerning the ethnolinguistic background (Lazaridis et al., 2014). Our results show that the Thai-HO group is quite similar to the CentralThai TK group (Figures 2-4; Supplementary Figure 10E; Supplementary Table 2), thus providing additional context for this group.

### South Asian Admixture

The South Asian (SA)-like signal in C_TK and S_TK groups is also one of the facilitating factors that enhance their differentiation from N_TK and NE_TK groups (Figures 2-4; Supplementary Figure 13). SA-related ancestry is also detected in the Mon and SouthernThai_AN (Figures 2-4; Supplementary Figure 13). SA admixture analyses indicated that the SA contribution to all Indian-related Thai groups is as a minor source (∼25%) while the main contribution comes from AA-related sources (Figure 6A). Although the CentralThai and SouthernThai_TK speak TK languages, and SouthernThai_AN speak an AN language, their genetic backgrounds are similar to AA groups (Figures 5B and 6A; Supplementary Figure 18), suggesting cultural diffusion to or admixture with AA groups. For the CentralThai, our previous mtDNA results showed admixture between Mon and CentralThai people, while the MSY results showed that the CentralThai were influenced by cultural diffusion from the Mon (Kutanan et al., 2018b, 2019). The SouthernThai_TK are genetically related to both the Mon and CentralThai (Figures 2 and 6C; Supplementary Figures 5, 16, and 18), consistent with historical evidence indicating that there were movements from the central region to the south during the Ayutthaya Period (during 1350-1767 A.D.) (Baker and Phongpaichit, 2017). Also living in the southern region, the SouthernThai_AN not only has SA-related ancestry, but it is also genetically distinct from AN-speaking groups from Taiwan (Ami and Atayal) and ISEA (Figure 2; Supplementary Figure 16). Similar to other SA-related groups, the SouthernThai_AN are more related to AA-speaking Cambodian and Khmu_Katu_AA groups in the PCA (Figure 2) and in the qpGraph received ancestry from a N_Indian ancestor (∼27%) and an admixed ancestor with Cambodian (∼90%) and Atayal (10%) ancestry (Figure 5B). This pattern is in agreement with the AN groups from Vietnam (Liu et al., 2020); our results support the MSEA origin of the SouthernThai_AN group, via cultural diffusion involving local AA groups.

There is archaeological evidence of frequent early prehistorical contacts between India and present-day Thailand (and Cambodia) during the Iron Age that brought exotic goods as well as ideas rooted in Buddhist and Hindu religions (Higham and Thodsarat, 2012). This could result in some Indian admixture in the local AA groups who then subsequently changed languages as a result of admixture or cultural diffusion involving arriving TK/AN groups. However, the dating of the Indian admixture in the Thai groups is more recent, ∼500-750 ya (Figure 6B; Supplementary Figure 17), which fits with the Ayutthaya Period (Baker and Phongpaichit, 2017). During the 16^th^ to 17^th^ century A.D., Siam (the former name for what is now the kingdom of Thailand) had maritime connections with westward trade dominated by Persians, Indians, Chinese and other nationalities who sailed from various Indian ports via the Melaka Straits or passed via Burmese ports to Ayutthaya (Baker and Phongpaichit, 2017; Ruangsilp and Wibulsilp, 2017). Trading and political connections – Indian Muslims served in administration (Chularatana, 2007) – would have facilitated admixture from South Asian to central Thai people (probably related to the Mon) during the Ayutthaya Period. As mentioned previously, this is also the time period of historical movements from the central region to the south, which could immediately bring the SA admixture to southern Thais (TK and AN). Alternatively, many ports in southern Thailand were also part of the international trade network, so the South Asian admixture in the southern Thais (TK and AN) probably also reflects this process. Europeans, e.g. Portuguese, were also an important part of this transnational network (Baker and Phongpaichit, 2017), but our results do not indicate any European genetic influence (Figures 2C and 6C; Supplementary Figure 5). Finally, a single-pulse admixture is inferred by GLOBETROTTER, which is supported by the admixture graph (Figure 6C; Supplementary Figure 18). Although this suggests that we have found a strong SA admixture signal from AA genetically related groups during the Ayutthaya Period, we cannot rule out the possibility of extensive and continuous interaction between South Asian and Mainland Southeast Asian in the past. More ancient DNA data from this region could provide further insights into this SA-MSEA interaction as well as the historical relationships among AA, TK, and AN groups in MSEA.

## Conclusions

We generated and analysed an extensive and intensive genome-wide SNP dataset from 36 ethnolinguistic groups from Thailand and Laos encompassing all five language families in MSEA, i.e. TK, AA, ST, HM and AN languages. We observed fine-scale genetic structure within each language family; interactions between AA and TK speakers are the principal factor influencing the population structure of the major TK speaking groups in each region. Interactions with South Asians also is evident in the genetic profiles of the Monic_AA, Central and Southern TK, and SouthernThai_AN groups. We also find genetic differences among ethnolinguistic groups within the ST and HM families, as well as among the hill tribes, that reflect different levels of contact with other groups. We observed genetic differentiation of the Thai and Taiwanese AN groups; genetic interactions between AN and AA groups in Thailand probably reflect cultural diffusion. Although our analyses provide the first detailed insights into the genetic history of Thai/Lao groups, further studies that include diverse modern groups from other MSEA countries, and more ancient samples, will provide even more insights into the demographic history of MSEA. In 2019, the Genomics Thailand Initiative was launched by the Thai government, with the goal of sequencing the genomes of 50,000 Thai people to enable precision medicine, and the project is ongoing. Our insights into the genetic structure of Thai/Lao ethnolinguistic groups should prove beneficial for selecting populations to include in such whole genome sequence and other biomedical studies.

## Material and Methods

### Sample preparation and quality control

Genomic DNA samples were from our previous studies (Kutanan et al., 2017; 2018; 2019) (Figure 1), with the exception of newly-collected samples from southern Thailand (SouthernThai_TK and SouthernThai_AN). In our previous studies, we interviewed all potential donors to screen for volunteers unrelated for at least two generations. We then collected blood, buccal or saliva samples with informed consent, which specified that their biological samples will also be stored for further anthropological genetic studies. For the present study, we used the same criteria as in the previous studies to recruit prospective donors from southern Thailand. Buccal samples were collected with written informed consent, and we extracted DNA using the Gentra Puregene Buccal Cell Kit (Qiagen, Germany) according to the manufacturer’s directions. Ethical approval for this study was granted by Khon Kaen University and by the Ethics Commission of the University of Leipzig Medical Faculty.

Genotyping was carried out using the Affymetrix Axiom Genome-Wide Human Origins array (Patterson et al., 2012); primary screening with the Affymetrix Genotyping Console v4.2 resulted in a total of 463 samples (genotype call rate >= 97%) genotyped for 596,085 loci on the hg19 version of the human reference genome coordinates.

We used PLINK version 1.90b5.2 (Purcell et al., 2007) to exclude loci and individuals with more than 5% missing data and also exclude mtDNA and sex chromosome loci. We further excluded loci which did not pass the Hardy-Weinberg equilibrium test (*p* value less than 0.00005), or had more than 50% missing data, within any population. We checked individual relatedness using KING (Manichaikul et al., 2010) implemented in PLINK version 2.0 (https://www.cog-genomics.org/plink/2.0/) and excluded one individual from each pair of individuals with 1^st^ degree kinship. There are in total 452 Thai/Lao individuals with 533,705 loci after these quality control measures (Supplementary Table 1).

We merged our data with data generated using the same array from modern populations from South Asia, East Asia and outgroup populations (the African Mbuti and European French) (Reich et al., 2011; Patterson et al., 2012; Lazaridis et al., 2014; Qin and Stoneking, 2015; Lipson et al., 2018) using mergeit in EIGENSOFT version 7.2.1 with default settings (Patterson et al., 2006). The data on ancient samples from previous studies (Lipson et al., 2018; McColl et al., 2018) were retrieved with all information included and their alleles were obtained through pseudo-haploid strategies. We excluded ancient samples with less than 15,000 informative loci; the number of loci after data merging is 370,732.

### Population structure analyses

For population structure analyses, PLINK version 1.90b5.2 was used to perform pruning for linkage disequilibrium, excluding one variant from pairs with r^2^ > 0.4 within windows of 200 variants and a step size of 25 variants, leaving in total 158,772 loci (153,191 loci when Mbuti and French are excluded). The Principle Component Analysis (PCA) was performed using smartpca from EIGENSOFT with the “lsqproject” and “autoshrink” options, with Mbuti and French excluded to focus on the structure among Asians. Three samples were identified as outliers based on the first 4 PCs and were removed (Supplementary Figure 2). The heatmap of additional PCs was visualized using the pheatmap package in R version 3.6.0. The clustering program ADMIXTURE version 1.3.0 (Alexander et al., 2009) was run from *K* = 2 to *K* = 15 with 100 replicates for each *K* and with random seeds with the -P option. The ancient samples and highly drifted modern populations (Onge, Mlabri, and Mamanwa) were projected in the PCA and ADMIXTURE analyses. PONG version 1.4.7 (Behr et al., 2016) was used to visualize the top 20 highest likelihood ADMIXTURE replicates for the major mode at each K.

### Allele sharing analyses

To test admixture and excess ancestry sharing, we computed *f3* and *f4*-statistics from ADMIXTOOLS version 5.1 (Patterson et al., 2012) using admixr version 0.7.1 (Petr et al., 2019), with significance assessed through block jackknife resampling across the genome and using Mbuti as the outgroup. Additional *f4*-statistics were computed using French as the outgroup to avoid deep attraction to Africans if ancient samples were involved, and only transversions (3,090-53,870 SNPs depending on the quality of samples) were used to avoid potential noise from ancient DNA damage patterns. The heatmap visualization of *f3* profiles was obtained using the pheatmap package in R.

### Data phasing and haplotype sharing analyses

To analyse haplotype sharing, we begin with data phasing; SHAPEIT version 4.1.3 (Delaneau et al., 2019) was used to phase the modern samples, with East Asian (without the Kinh Vietnamese merged in our dataset) and South Asian populations as a reference panel, and the recombination map from the 1000 Genomes Phase3 (Genomes Project et al., 2015). To prepare the reference panel, we extracted the East and South Asian individuals as well as the overlapping sites with our data for each chromosome from the 1000 Genomes Phase3 data using bcftools version 1.4 (http://samtools.github.io/bcftools/). The phasing accuracy of SHAPEIT4 can be enhanced by increasing the number of conditioning neighbors in the Positional Burrows–Wheeler Transform (PBWT) on which haplotype estimation is based (Delaneau et al., 2019). We ran phasing with the options --pbwt-depth 8 for 8 conditioning neighbors and left other parameters as default.

We then ran ChromoPainter version 2 (Lawson et al., 2012) on the phased data set to begin the haplotype sharing investigation, with sample sizes for each population randomly down-sampled to 4 and 8. The former was used for 10 iterations of the EM (expectation maximization) process to estimate the switch rate and global mutation probability, while the latter was for the chromosomal painting process with the estimated switch and global mutation rates, which then gave the output for downstream analyses. We first attempted to paint the chromosomes of each individual, using all of the modern Asian samples as both donors and recipients via the -a argument. The EM estimation of switch rate and global mutation probability were ∼623.09 and ∼0.0013, respectively, which were then used as the starting values for these parameters for all donors in the painting process. To minimize the effect of genetic drift in the Thai/Lao groups, we also performed another run using all the modern Asian samples except for those sampled in this study as both donors and recipients; samples from this study were used only as recipients. The EM estimation of switch rate and global mutation probability for this analysis were ∼764.56 and ∼0.0011, respectively. The heatmap results were generated using the pheatmap package in R.

To identify shared IBD blocks between each pair of individuals and homozygous-by-descent (HBD) blocks within each individual, we used refinedIBD (Browning and Browning, 2013). Both identified IBD and HBD blocks are considered as IBD blocks in our analyses, which is analogous to pairwise shared coalescence (PSC) segments in a previous study (Al-Asadi et al., 2019). The IBD blocks within a 0.6 cM gap were merged using the program merge-ibd-segments from BEAGLE utilities (Browning and Browning, 2007; Browning et al., 2018), allowing only 1 inconsistent genotype between the gap and block regions. These results were used to generate four datasets based on the identified IBD blocks lengths: 1 to 5 cM, 5 to 10 cM, over 10 cM, and at least 2 cM. We used the first three datasets for analysis of the IBD sharing between populations by network visualization in different time periods (Ralph and Coop, 2013; Al-Asadi et al., 2019), while the last one was used to analyse overall IBD sharing between populations by heatmap and IBD sharing within each individual population (Browning and Browning, 2015; Browning et al., 2018). In each dataset, we summed up the total number and length of IBD blocks for each individual pair and calculated the population median and mean. The pairs with at least 10 cM average summed length (4 cM for the range of 1 to 5 cM) of shared blocks were kept to reduce noise and false positives in network visualization. The IBDNe software (Browning and Browning, 2015; Browning et al., 2018) was employed to estimate effective population size changes over time with the following conditions as suggested previously (Browning and Browning, 2015): shared blocks of at least 2cM within each population, and estimated population size numbers inferred within the past 50 generations only, as previously suggested for SNP array data (Browning and Browning, 2015). A generation time of 30 years (Fenner, 2005) was used to convert generations to years.

### Admixture source and date inferences

For the populations with apparent Indian admixture, we ran GLOBETROTTER (Hellenthal et al., 2014) using the ChromoPainter results with only Thai/Lao samples in this study as recipients and all the donors as surrogates. We first tested the certainty and potential waves of admixture events, and then estimated the major and minor sources as well as the dates of admixture. The distributions of admixture dates were accessed through 100 bootstraps. We also dated admixture events with ALDER (Loh et al., 2013) using the populations identified as the major (Kinh) and minor (Gujarati) sources in the GLOBETROTTER analysis as the two sources used to date the admixture in the ALDER analysis. However, we could not get an acceptable fit of the LD decay curves between Kinh and all the tested groups, so we present the dates inferred using Gujarati as a single source instead. Again, genetic map information was retrieved from 1000 Genomes Phase3 data (Genomes Project et al., 2015).

### Admixture graph analyses

Using the pruned dataset (18,310 SNPs) of the Thai/Lao and other reference modern populations (based on ChromoPainter results) and ancient samples (with more than 130,000 overlapping SNPs, corresponding to < 65% missing data), TreeMix version 1.12 (Pickrell and Pritchard, 2012) was used to construct a maximum-likelihood tree in order to reveal population relationships and migration among five ancient samples (Ho-PhaFaen, N-GuaChacCave, N-TamPaLing, IA-LongLongRak and Hi-Kinabatagan), Thai/Lao modern populations, and selected reference modern populations, i.e. the African Mbuti (used as outgroup), European French, Indo-European-speaking Indian groups (Gujarati, Brahmin Tiwari, and Lodhi), Andamanese Onge, and East Asian groups from the five different language families (AA-speaking Cambodian, TK-speaking Dai, AN-speaking Atayal, ST-speaking Naxi and HM-speaking Miao). The Indo-European-speaking Indian groups were together labelled as N_Indian as they are enriched for the “North Indian” ancestry component identified previously, whereas Onge are enriched for “South Indian” ancestry (Reich et al., 2009). Based on ChromoPainter results, the AA Thai groups were further grouped into Monic_AA (Mon), Khmu_Katu_AA (HtinMal, HtinPray, Mlabri, Khmu, So, and Bru) and Palaungic_AA (Lawa_Eastern, Lawa_Western, Palaung, and Blang); the TK Thai/Lao groups were grouped into N_TK (Khonmueang, Shan, Khuen, Lue, Phuan, and Yuan), NE_TK (Black Tai, LaoIsan, Phutai, Nyaw, Kalueang and Laotian), C_TK (CentralThai) and S_TK (SouthernThai_TK); the HmongNjua and HmongDaw were grouped into Hmong_HM; and the KarenPwo, KarenPadaung, and KarenSkaw were grouped into Karen_ST. We investigated 0 to 3 migration events using 10 independent runs and then selected the topology with the highest likelihood for further investigation. To model admixture graphs, we used AdmixtureBayes (Nielsen, 2018) to estimate the top 10 posterior admixture graphs for Thai/Lao groups from each language family and comparative modern populations (including the associated linguistic source groups, N_Indian group, and outgroup Mbuti), based on the covariance of the allele frequency profiles. We also performed an additional investigation of the potential South Asian genetic influence on some Thai groups (Mon, C_TK, S_TK, SouthernThai_AN), including Mbuti, French, N_Indian, Onge, and the associated linguistic source groups to disentangle potential East Asian vs. South Indian/Hoabihian (Onge) vs. North Indian (N_Indian) vs. European (French) ancestry. Each case study graph was inferred from an independent pruned dataset with 175,578-191,384 SNPs, depending on the number of groups/individuals. For each AdmixtureBayes run, a total of 300,000 MCMC steps were carried out, stopping the run if the summaries of effective sample size were all above 200. Finally, we used the estimated graphs from AdmixtureBayes as input for qpGraph from ADMIXTOOLS to test the goodness of fit of the graphs. Acceptable graphs have, by convention, an absolute value of the Z-score of the worst *f4* statistic less than 3. If none of the estimated graphs from AdmixtureBayes produced an acceptable graph, we removed populations based on the *f4* outliers output of qpGraph, used the option “-subnodes” in AdmixtureBayes, and ran qpGraph again. We iterated these procedures until we were able to find an acceptable graph. The qpGraph parameters are as follows: outpop: NULL, blgsize: 0.05, forcezmode: YES, diag: 0.0001, bigiter: 6, hires: YES, and lambdascale: 1.

## Acknowledgements

We thank all sample donors for making this work possible. We acknowledge Khamnikone Sipaseuth, Saksuriya Triyarach, Narongdech Mahasirikul, Supada Khonyoung and Dusit Boonmekam for assistance in collecting samples. We thank Roland Schröder for technical assistance. We thank Sandra Oliveira and Irina Pugach for helpful advice concerning computational analyses. This study was supported by the Max Planck Society and the Thailand Research Fund (RSA6180058).

## Data Availability

Data are made available upon receipt of a signed letter to the corresponding author confirming that the data will only be used in accordance with the restrictions of the informed consent, including the following: the data will not be transferred to anyone else; the data will be used for genetic/anthropological studies but not for any commercial purposes or no attempt to identify any of the sample donors.

