## Supplementary Figures for "Reconstructing the human genetic history of mainland Southeast Asia: insights from genome-wide data from Thailand and Laos"

There are 18 supplementary figures and 3 supplementary tables in this manuscript.

**Supplementary Figure 1** Sample information map for the comparative data used in this study. (A) Modern samples colored according to language family. (B) Ancient samples colored according to time period.

**Supplementary Figure 2** PCA plots of all modern individuals from the Thai/Lao and comparative Asian populations before outliers were removed, colored according to language family. (A) PC1 vs. PC2. (B) PC3 vs. PC4. The dataset consists of 845 individuals and 153,191 SNPs. The highly drifted modern populations (Onge, Mlabri, and Mamanwa) were projected in the PCA analysis. Individuals BST109, PUT101, and KR117 were removed from downstream analyses as PCA outliers.

**Supplementary Figure 3** Plot of PC1 vs. PC2, with modern individuals colored according to (A) country and (B) language family; ancient individuals are labeled in black. The dataset consists of 875 individuals and 153,191 SNPs. The highly drifted modern populations (Onge, Mlabri, and Mamanwa) and ancient samples were projected in the PCA analysis.

**Supplementary Figure 4** Cross validation errors of ADMIXTURE runs for $K$= 2 to $K$ = 15, based on 100 runs for each $K$ value.

**Supplementary Figure 5** ADMIXTURE results of modern and ancient data for $K$ = 2-15. Each individual is represented by a thin vertical line, which is partitioned into $K$ colored segments that represent the individual's estimated membership fraction for each of the $K$ ancestry components. Populations are separated by black lines. The highly drifted modern populations (Onge, Mlabri, and Mamanwa) and ancient samples were projected in the ADMIXTURE analyses. The three colored bars at the top of the plot indicate the country (top), language family (middle) and subgroup (bottom) for each sample, according to the key at the bottom.

**Supplementary Figure 6** Heat map of shared haplotype segments estimated via Chromopainter. The recipients (Thai/Lao groups) are labelled on the X-axis and the donors are indicated on the Y-axis. The heatmap is scaled with the average length in centimorgans of the summed painted chromosomal chunks of the recipient individuals from the donor individuals.

**Supplementary Figure 7** IBD sharing within each Thai/Lao group. (A) with Mlabri and (B) without Mlabri. The Y axis and X axis indicate the mean number and summed length of IBD blocks within each group, respectively.

**Supplementary Figure 8** Mean number of shared blocks between populations. (A) Network visualizations of the mean number of shared IBD blocks focusing on the sharing involving Thai/Lao populations within MSEA, with identified IBD blocks in the range of 1 to 5 cM, 5 to 10 cM, and over 10 cM, with the mean length of summed IBD block at least 4 cM, 10 cM, and 10 cM, respectively. (B) Network visualizations of the mean number of shared IBD blocks focusing on the sharing involving Thai/Lao populations within East Asia with identified IBD blocks in the range of 1 to 5 cM and the mean length of summed IBD blocks at least 4 cM. Each circle stands for a population, colored according to language family, and each edge (colored according to the scale) indicates the IBD sharing between populations.

**Supplementary Figure 9** Effective population size change over time for each Thai/Lao group over the past 50 generations, estimated from the within-group shared IBD. Solid lines are the effective population size change; 95% confidence intervals are shaded in grey.

**Supplementary Figure 10** *f4* statistics comparing two Thai/Lao populations from the same language family (LF) with another Thai/Lao population from a different LF. Z-scores are for *f4* (X, Y; Z, Mbuti), where X is a selected Thai/Lao population in LF1, Y is another population also in LF1, and Z is population not in LF1. Different symbols denote different populations for Y. The dots are colored according to the LF of Z. The vertical red dashed lines denote +3/-3 while the solid line denotes 0. We show here representative examples of the results, where the selected population X is (A) IuMien in HM, (B) KarenPadaung in ST, (C) Palaung in AA, (D) Laotian in TK, and (E) CentralThai in TK. All of the *f4* statistics comparing two Thai/Lao populations from the same LF to another Thai/Lao population from a different LF are in Supplementary Table 2.

**Supplementary Figure 11** *f4* statistics comparing comparing Thai/Lao populations from two different subgroups (SGs) of the same LF. Z-scores are for *f4* (X, Y; Z, Mbuti), where X is a selected Thai/Lao population in SG1, Y is another population also in SG1, and Z is a population not in SG1. Different symbols denote different populations for Y. The dots are colored according to the SG of Z. The vertical grey dashed lines denote +3/-3 while the solid line denotes 0. We show here representative examples of the results, where the selected population X is (A) Laotian in NE_TK, (B) LaoIsan in NE_TK, (C) Khonmueang in N_TK, (D) Palaung in Palaungic_AA, and (E) Soa in Khmu_Katu_AA. All of the *f4* statistics comparing two Thai/Lao populations from the same SG to other Thai/Lao populations are in Supplementary Table 3.

**Supplementary Figure 12** *f4* statistics comparing Thai/Lao populations to representative East Asian populations and Han Chinese. Z-scores are for *f4* (W, Han; Y, Mbuti), where W is the selected East Asian population (panel labels) and Y is the Thai/Lao population (label on the Y axis). The vertical grey lines denote 0. The panels, dots, and error bars are colored according to language family. Empty circles denote nonsignificant Z-scores (|Z|=<3) and solid circles denote significant Z-scores (|Z|>3). The colored bar denotes subgroups of Thai/Lao groups, according to the key on the right.

**Supplementary Figure 13** *f4* statistics comparing Thai/Lao populations to Indian populations. Z-scores are for *f4* (Thai/Lao population, Han; Indian population, Mbuti); each panel shows the results for the Indian population (panel label) compared to all Thai/Lao groups (Y-axis labels). The vertical grey lines denote 0. The dots, and error bars are colored according to language family. Empty circles denote nonsignificant Z-scores (|Z|=<3) and solid circles denote significant Z-scores (|Z|>3). The colored bar to the right of the group names denotes linguistic/geographic subgroups of Thai/Lao groups, according to the key on the right.

**Supplementary Figure 14** *f4* statistics comparing Thai/Lao populations to ancient samples. Z-scores are for f4 (ancient samples, Han; Thai/Lao populations, French). The vertical grey lines denote 0. The dots and error bars are colored according to language family. Empty circles denote nonsignificant Z-scores (|Z|=<3) and solid circles denote significant Z-scores (|Z|>3). The colored bar denotes subgroups of Thai/Lao groups, according to the key on the right.

**Supplementary Figure 15** TreeMix results for the Thai/Lao groups and representative modern populations and ancient samples, (A) The maximum likelihood tree without migration and the corresponding residual plot and (B) the maximum likelihood tree with three migration events and the corresponding residual plot. Thai/Lao groups are colored according to language family.

**Supplementary Figure 16** Alternative admixture graph for the TK groups with EA source populations included (worst-fitting Z = -7.037). The node r denotes the root. White nodes denote backbone populations. Backbone population labels and Thai nodes are colored according to language family. Dashed arrows represent admixture edges, while solid arrows are drift edges reported in units of FST×1,000.

**Supplementary Figure 17** Estimated dates for the SA-related admixture in four putative SA-influenced Thai groups (SouthernThai_AN, SouthernThai_TK, CentralThai and Mon). ALDER estimated dates are in red, using Gujarati as a single source; GLOBETROTTER estimated dates are in grey. Note that ALDER dating failed for the Yuan.

**Supplementary Figure 18** Alternative admixture graph for the SA-related Thai groups with EA source populations included and Mbuti as outgroup (worst-fitting Z = -4.110). The node r denotes the root. White nodes denote backbone populations. Backbone population labels and Thai nodes are colored according to language family. Dashed arrows represent admixture edges, while solid arrows are drift edges reported in units of FST×1,000. The star-shaped node denotes the N_Indian-related source contributing to all of the SA-related Thai groups. The AA-related lineage is highlighted in purple.

**Supplementary Table 1** General information concerning populations and samples studied. (excel file)

**Supplementary Table 2** The result of *f4*-statistics of the form *f4* (group 1, group 2; group 3, Outgroup), where Outgroup = Mbuti, group 1 and group 2 are groups from the same language family, and group 3 is from a different language family. BABA is the number of SNPs where groups 1 and 3 share the same allele and groups 2 and 4 share the alternative allele; ABBA is the number of SNPs where groups 1 and 4 share the same allele and groups 2 and 3 share the alternative allele. The total number of SNPs is 370,732 for each comparison. (excel file)

**Supplementary Table 3** The result of *f4*-statistics of the form *f4* (group 1, group 2; group 3, Outgroup), where Outgroup = Mbuti, group 1 and group 2 are groups from the same subgroup, and group 3 is from a different subgroup. BABA is the number of SNPs where groups 1 and 3 share the same allele and groups 2 and 4 share the alternative allele; ABBA is the number of SNPs where groups 1 and 4 share the same allele and groups 2 and 3 share the alternative allele. The total number of SNPs is 370,732 for each comparison. (excel file)

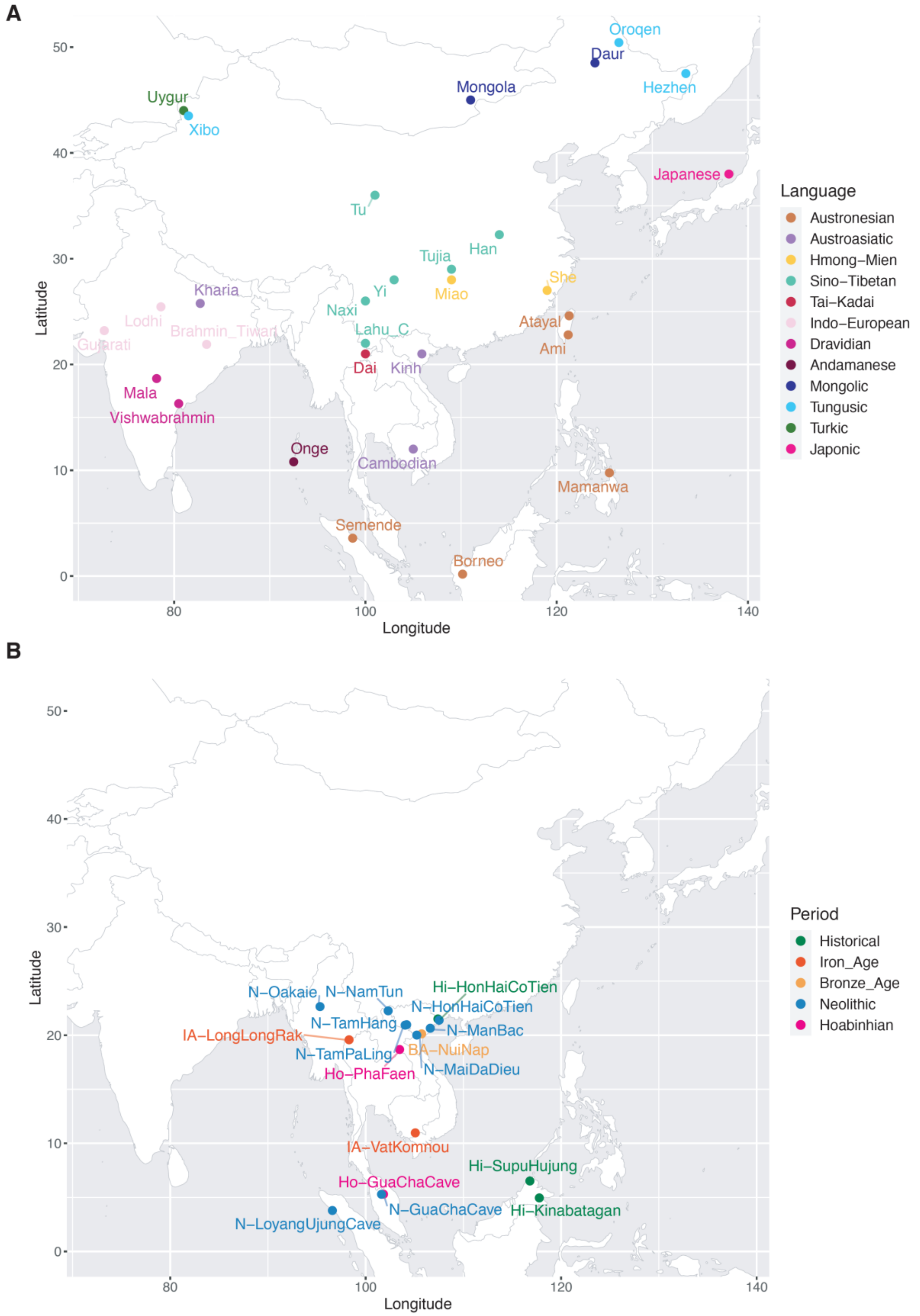


**Supplementary Figure 1** Sample information map for the comparative data used in this study. (A) Modern samples colored according to language family. (B) Ancient samples colored according to time period.

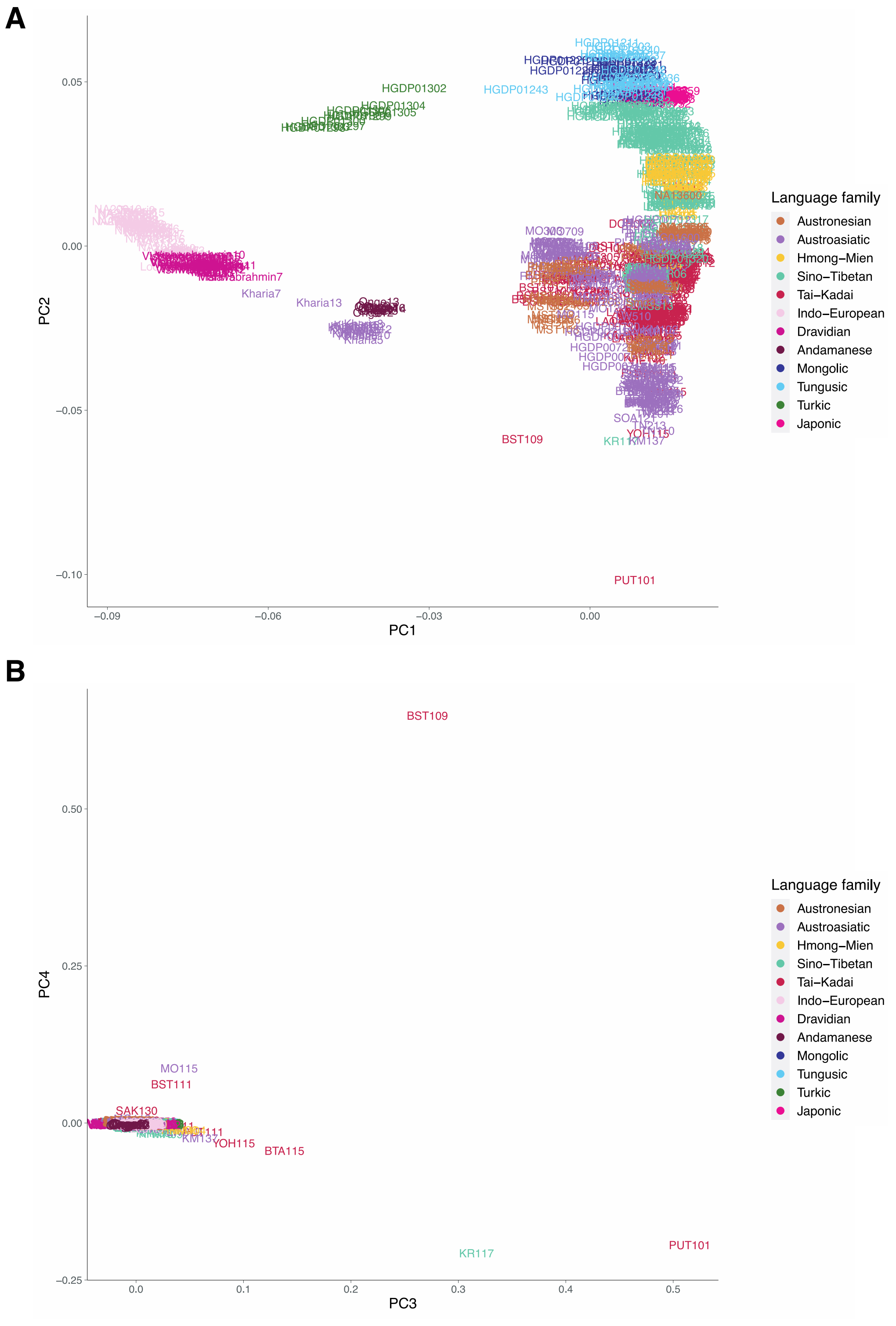


**Supplementary Figure 2** PCA plots of all modern individuals from the Thai/Lao and comparative Asian populations before outliers were removed, colored according to language family. (A) PC1 vs. PC2. (B) PC3 vs. PC4. The dataset consists of 845 individuals and 153,191 SNPs. The highly drifted modern populations (Onge, Mlabri, and Mamanwa) were projected in the PCA analysis. Individuals BST109, PUT101, and KR117 were removed from downstream analyses as PCA outliers.

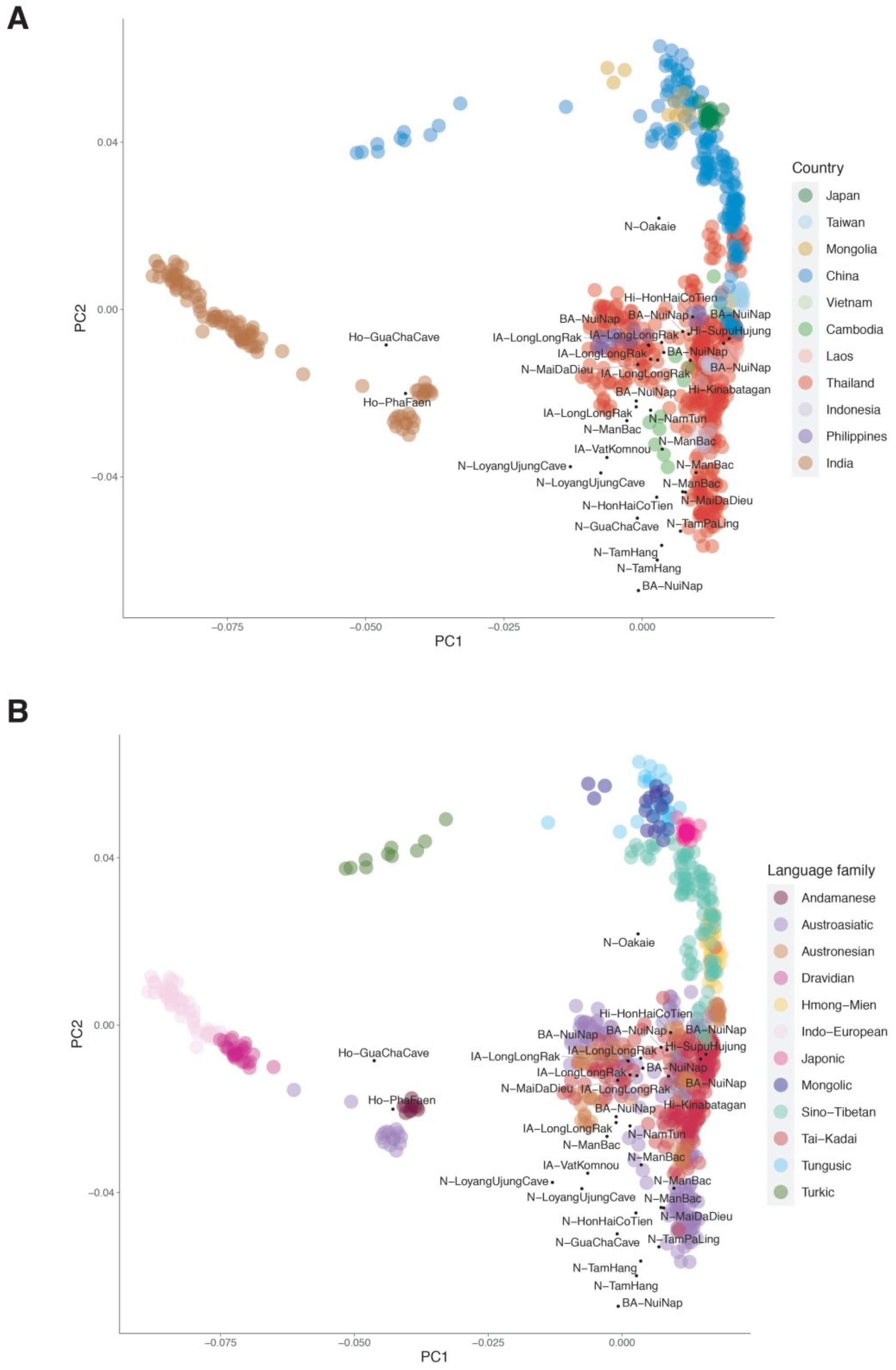


**Supplementary Figure 3** Plot of PC1 vs. PC2, with modern individuals colored according to (A) country and (B) language family; ancient individuals are labeled in black. The dataset consists of 875 individuals and 153,191 SNPs. The highly drifted modern populations (Onge, Mlabri, and Mamanwa) and ancient samples were projected in the PCA analysis.

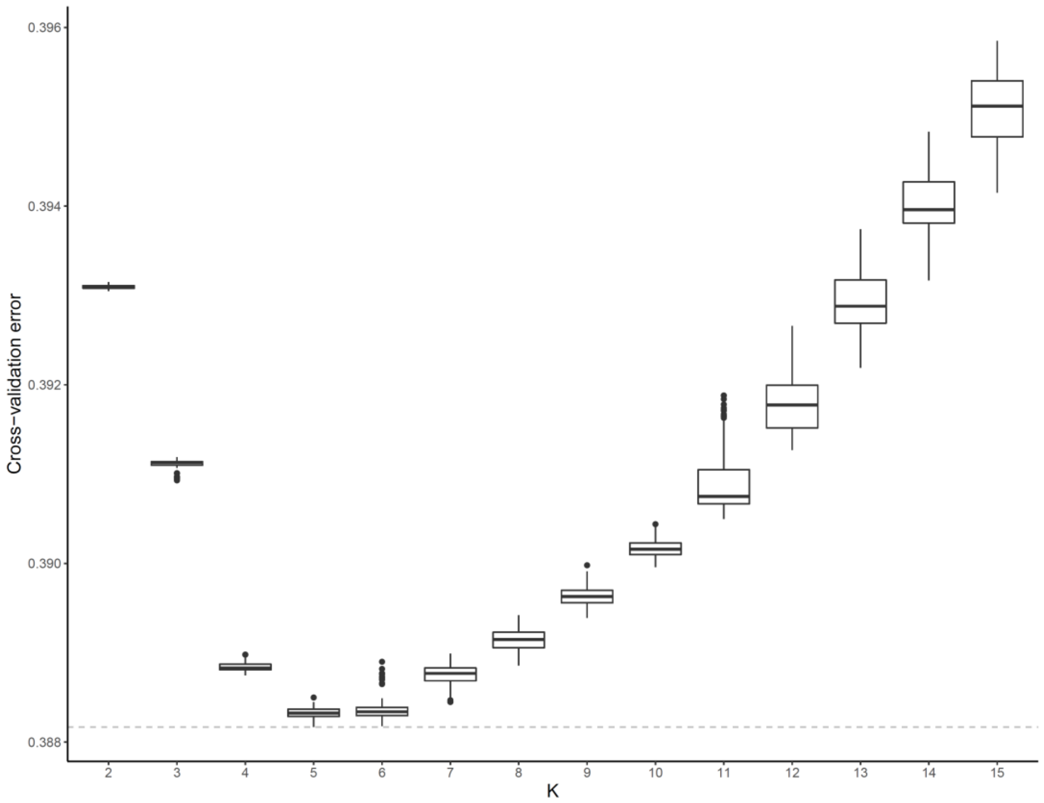


**Supplementary Figure 4** Cross validation errors of ADMIXTURE runs for *K*= 2 to *K* = 15, based on 100 runs for each *K* value.

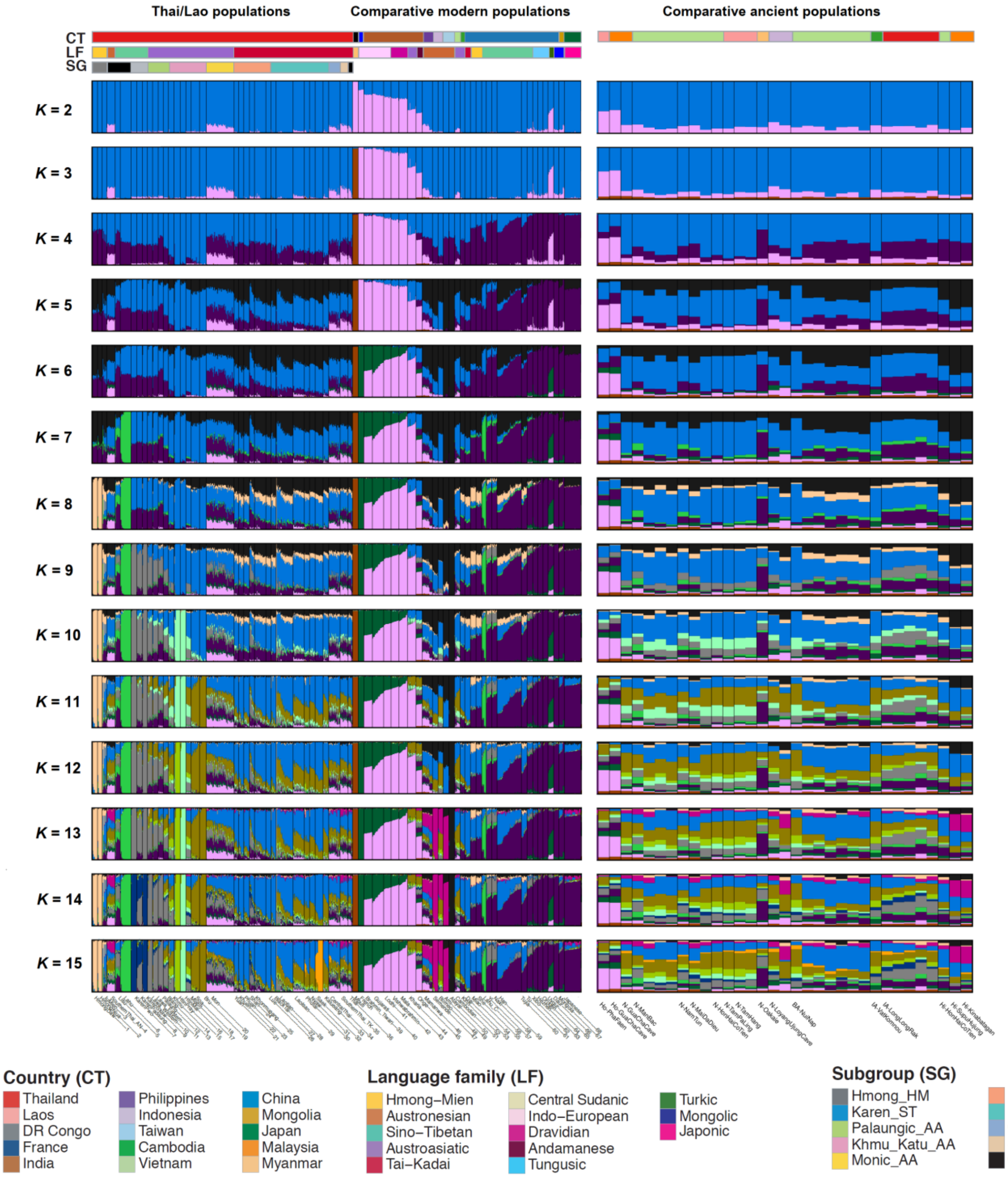


**Supplementary Figure 5** ADMIXTURE results of modern and ancient data for $K$ = 2-15. Each individual is represented by a thin vertical line, which is partitioned into $K$ colored segments that represent the individual's estimated membership fraction for each of the $K$ ancestry components. Populations are separated by black lines. The highly drifted modern populations (Onge, Mlabri, and Mamanwa) and ancient samples were projected in the ADMIXTURE analyses. The three colored bars at the top of the plot indicate the country (top), language family (middle) and subgroup (bottom) for each sample, according to the key at the bottom.

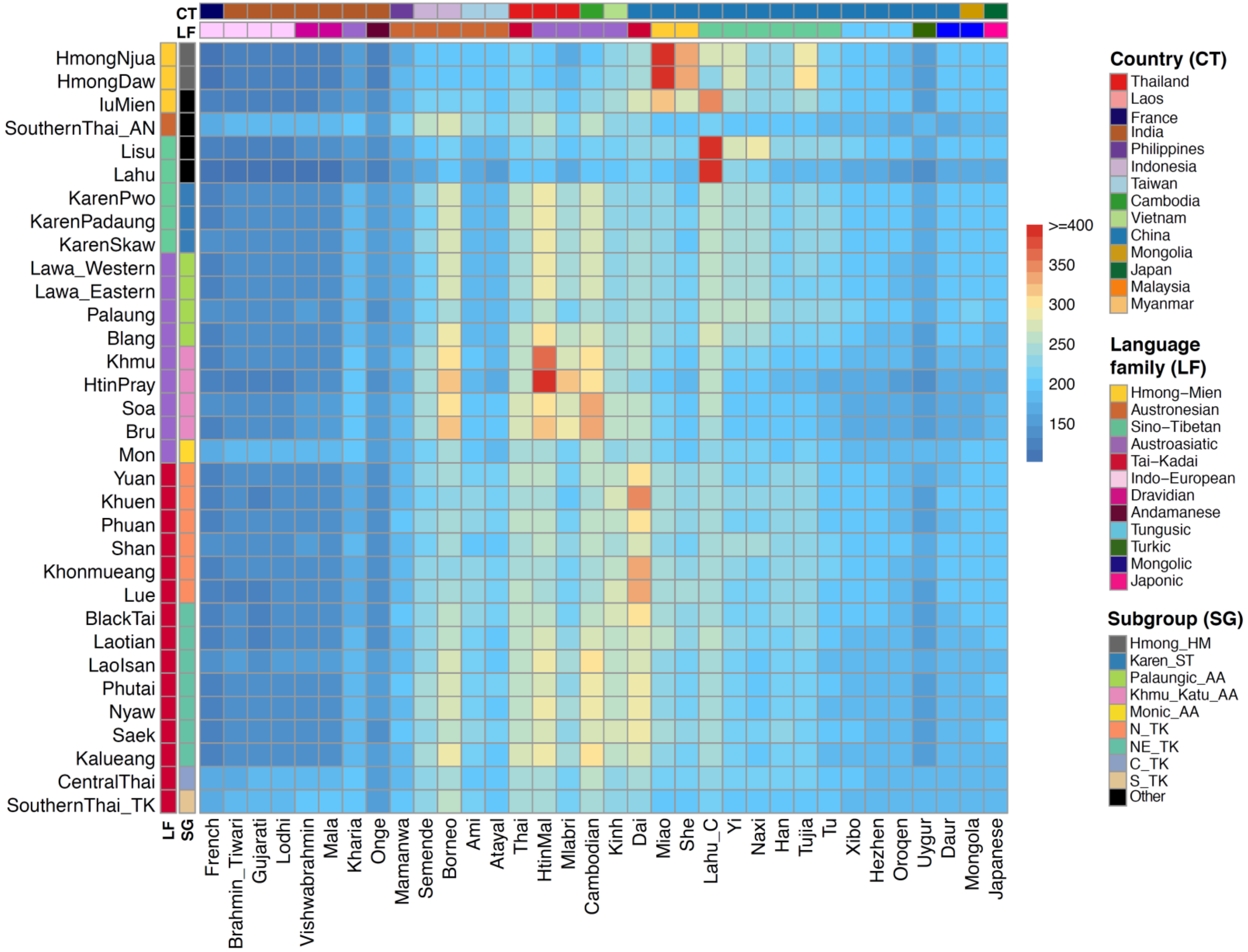


**Supplementary Figure 6** Heat map of shared haplotype segments estimated via Chromopainter. The recipients (Thai/Lao groups) are labelled on the X-axis and the donors are indicated on the Y-axis. The heatmap is scaled with the average length in centimorgans of the summed painted chromosomal chunks of the recipient individuals from the donor individuals.

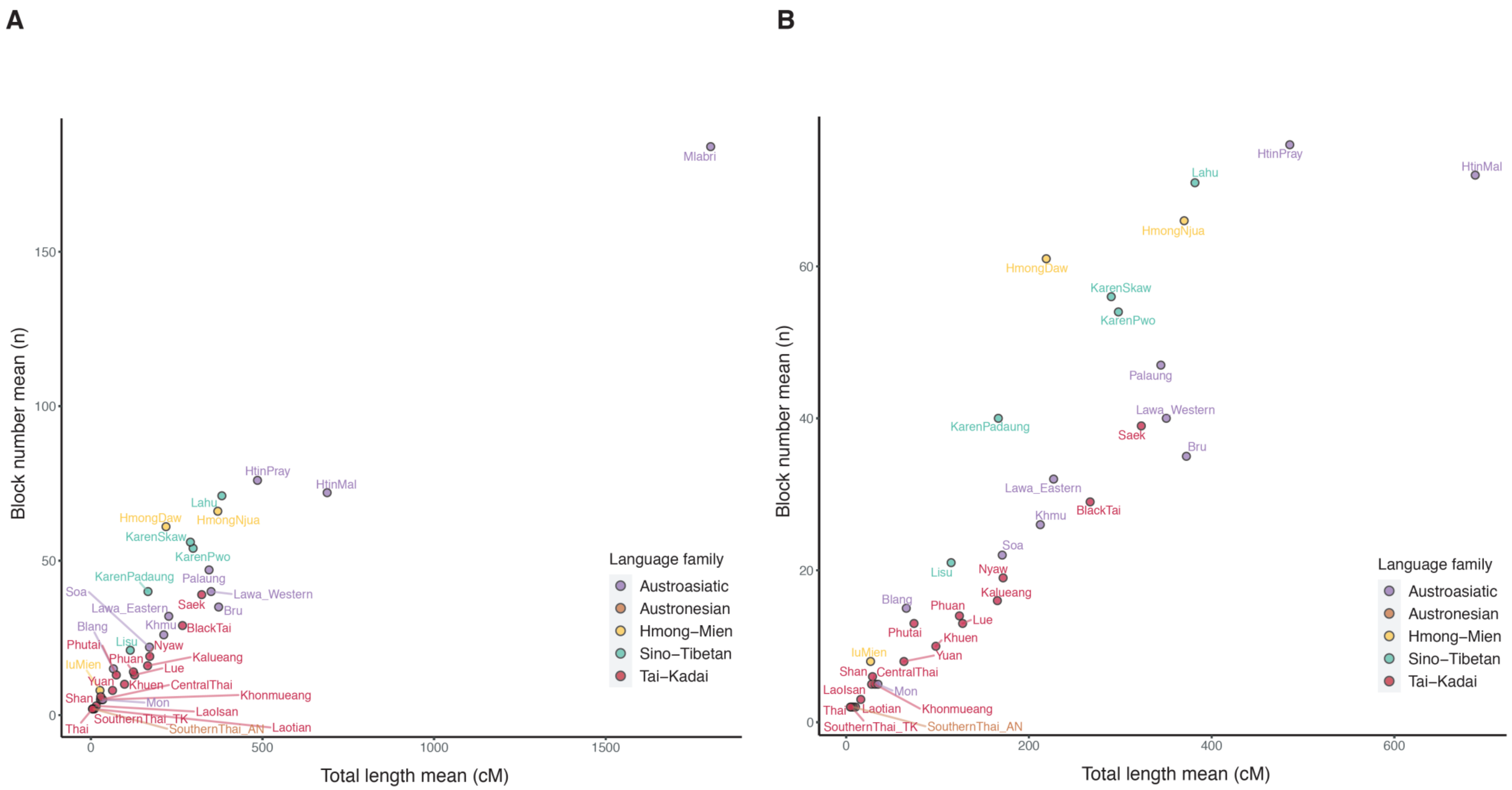


**Supplementary Figure 7** IBD sharing within each Thai/Lao group. (A) with Mlabri and (B) without Mlabri. The Y axis and X axis indicate the mean number and summed length of IBD blocks within each group, respectively.

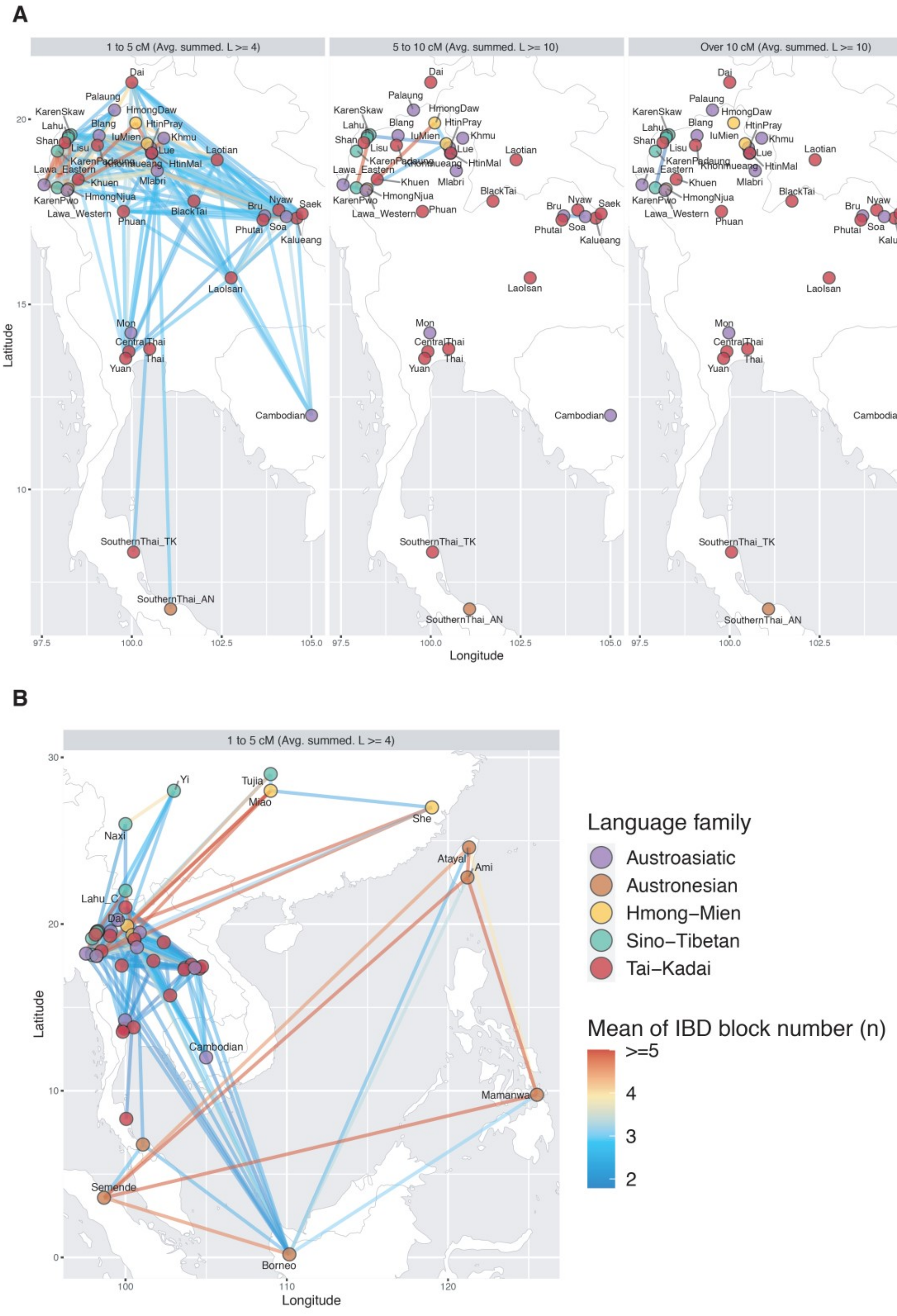


**Supplementary Figure 8** Mean number of shared blocks between populations. (A) Network visualizations of the mean number of shared IBD blocks focusing on the sharing involving Thai/Lao populations within MSEA, with identified IBD blocks in the range of 1 to 5 cM, 5 to 10 cM, and over 10 cM, with the mean length of summed IBD block at least 4 cM, 10 cM, and 10 cM, respectively. (B) Network visualizations of the mean number of shared IBD blocks focusing on the sharing involving Thai/Lao populations within East Asia with identified IBD blocks in the range of 1 to 5 cM and the mean length of summed IBD blocks at least 4 cM. Each circle stands for a population, colored according to language family, and each edge (colored according to the scale) indicates the IBD sharing between populations.

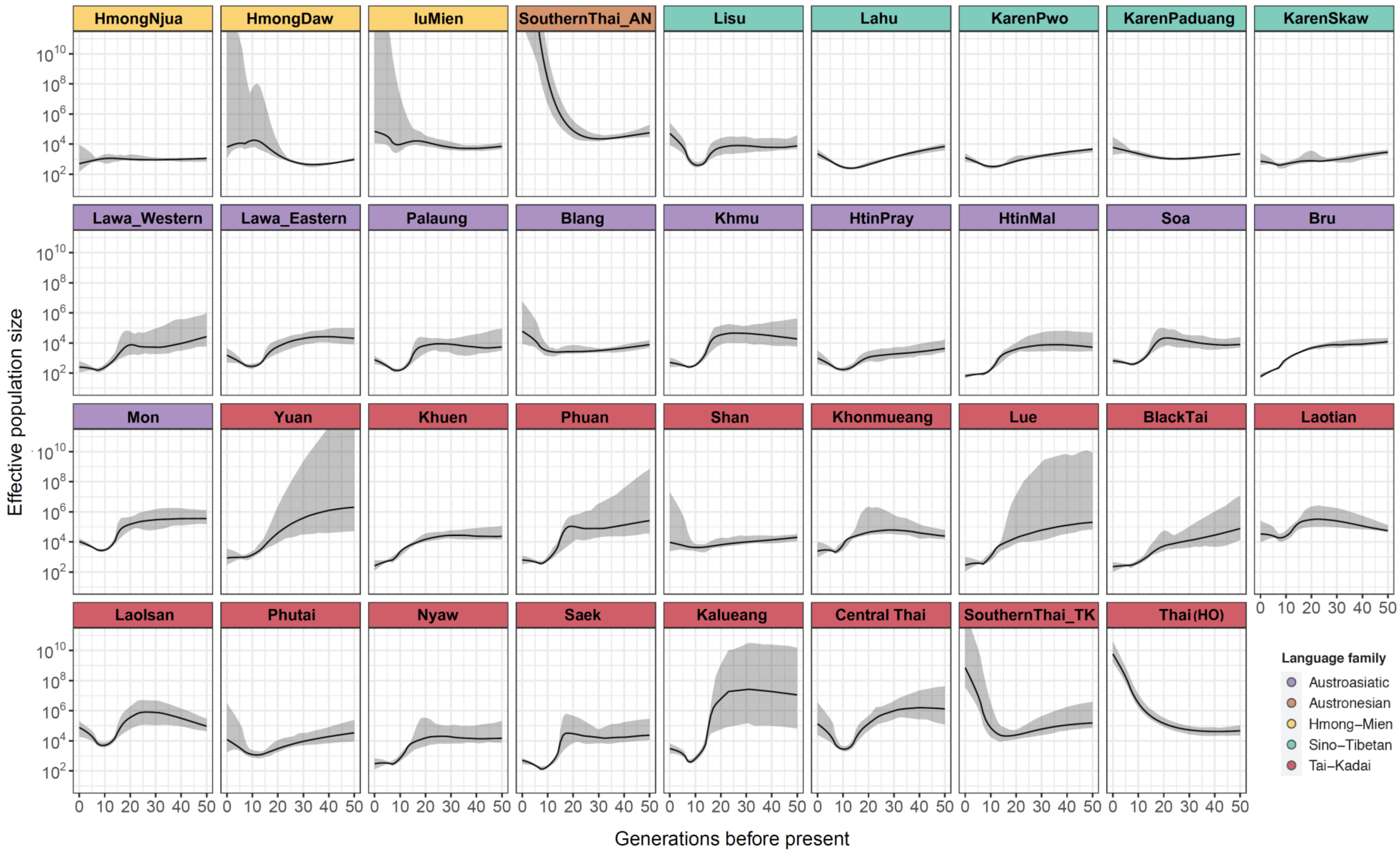


**Supplementary Figure 9** Effective population size change over time for each Thai/Lao group over the past 50 generations, estimated from the within-group shared IBD. Solid lines are the effective population size change; 95% confidence intervals are shaded in grey.

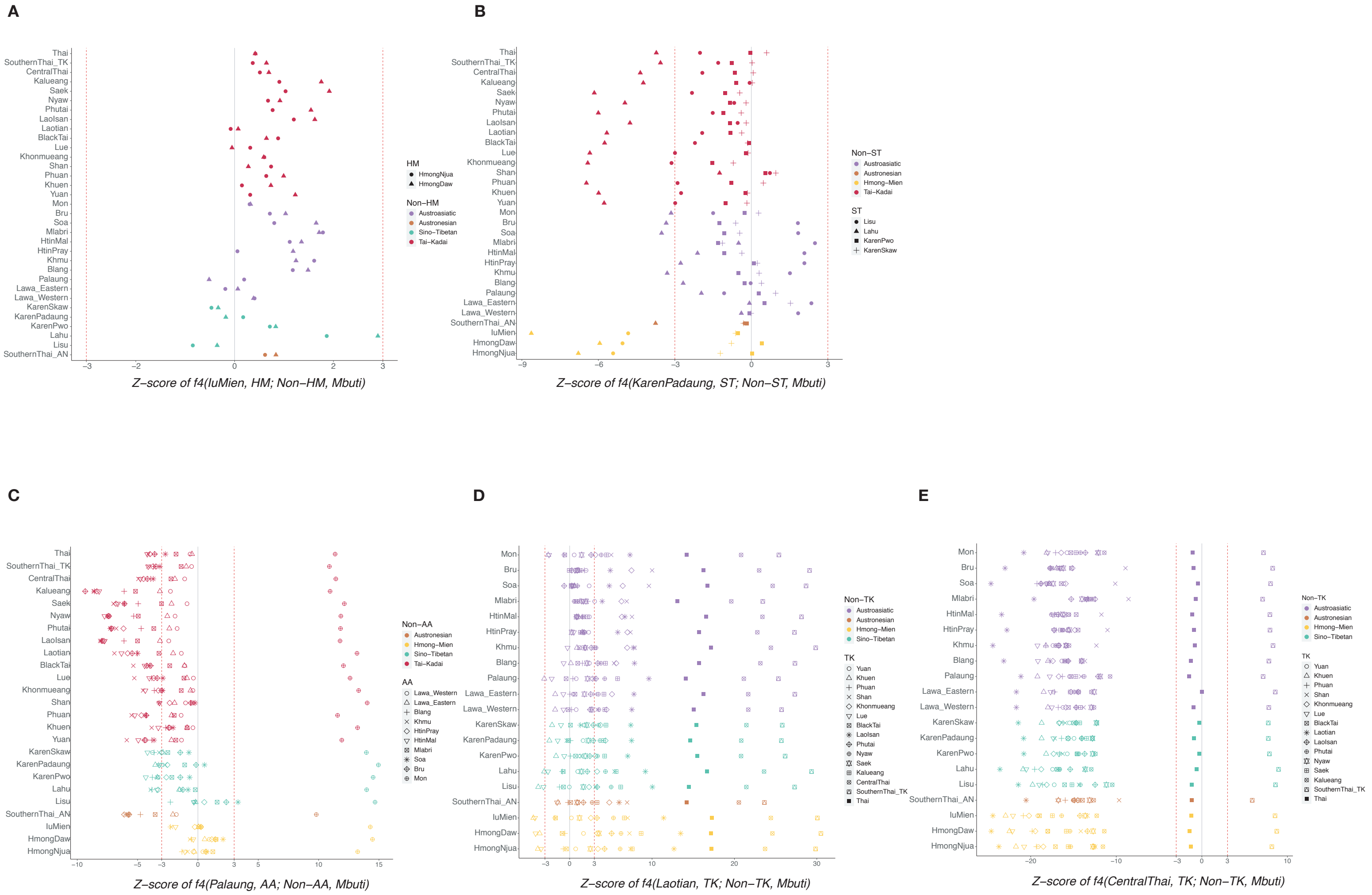

**Supplementary Figure 10** *f4* statistics comparing two Thai/Lao populations from the same language family (LF) with another Thai/Lao population from a different LF. Z-scores are for *f4* (X, Y; Z, Mbuti), where X is a selected Thai/Lao population in LF1, Y is another population also in LF1, and Z is population not in LF1. Different symbols denote different populations for Y. The dots are colored according to the LF of Z. The vertical red dashed lines denote +3/-3 while the solid line denotes 0. We show here representative examples of the results, where the selected population X is (A) IuMien in HM, (B) KarenPadaung in ST, (C) Palaung in AA, (D) Laotian in TK, and (E) CentralThai in TK. All of the *f4* statistics comparing two Thai/Lao populations from the same LF to another Thai/Lao population from a different LF are in Supplementary Table 2.

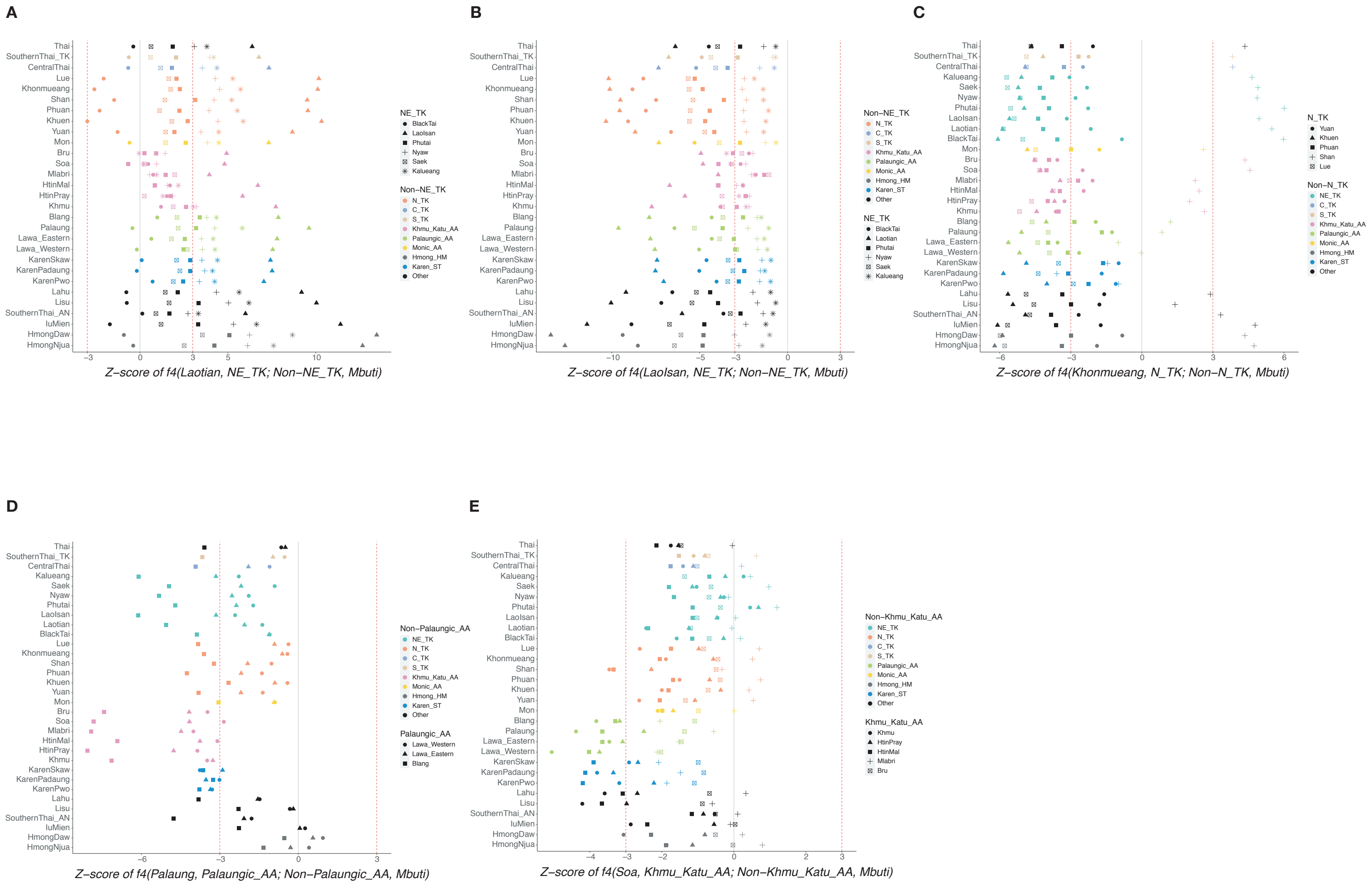


**Supplementary Figure 11** *f4* statistics comparing comparing Thai/Lao populations from two different subgroups (SGs) of the same LF. Z-scores are for *f4* (X, Y; Z, Mbuti), where X is a selected Thai/Lao population in SG1, Y is another population also in SG1, and Z is a population not in SG1. Different symbols denote different populations for Y. The dots are colored according to the SG of Z. The vertical grey dashed lines denote +3/-3 while the solid line denotes 0. We show here representative examples of the results, where the selected population X is (A) Laotian in NE_TK, (B) LaoIsan in NE_TK, (C) Khonmueang in N_TK, (D) Palaung in Palaungic_AA, and (E) Soa in Khmu_Katu_AA. All of the *f4* statistics comparing two Thai/Lao populations from the same SG to other Thai/Lao populations are in Supplementary Table 3.

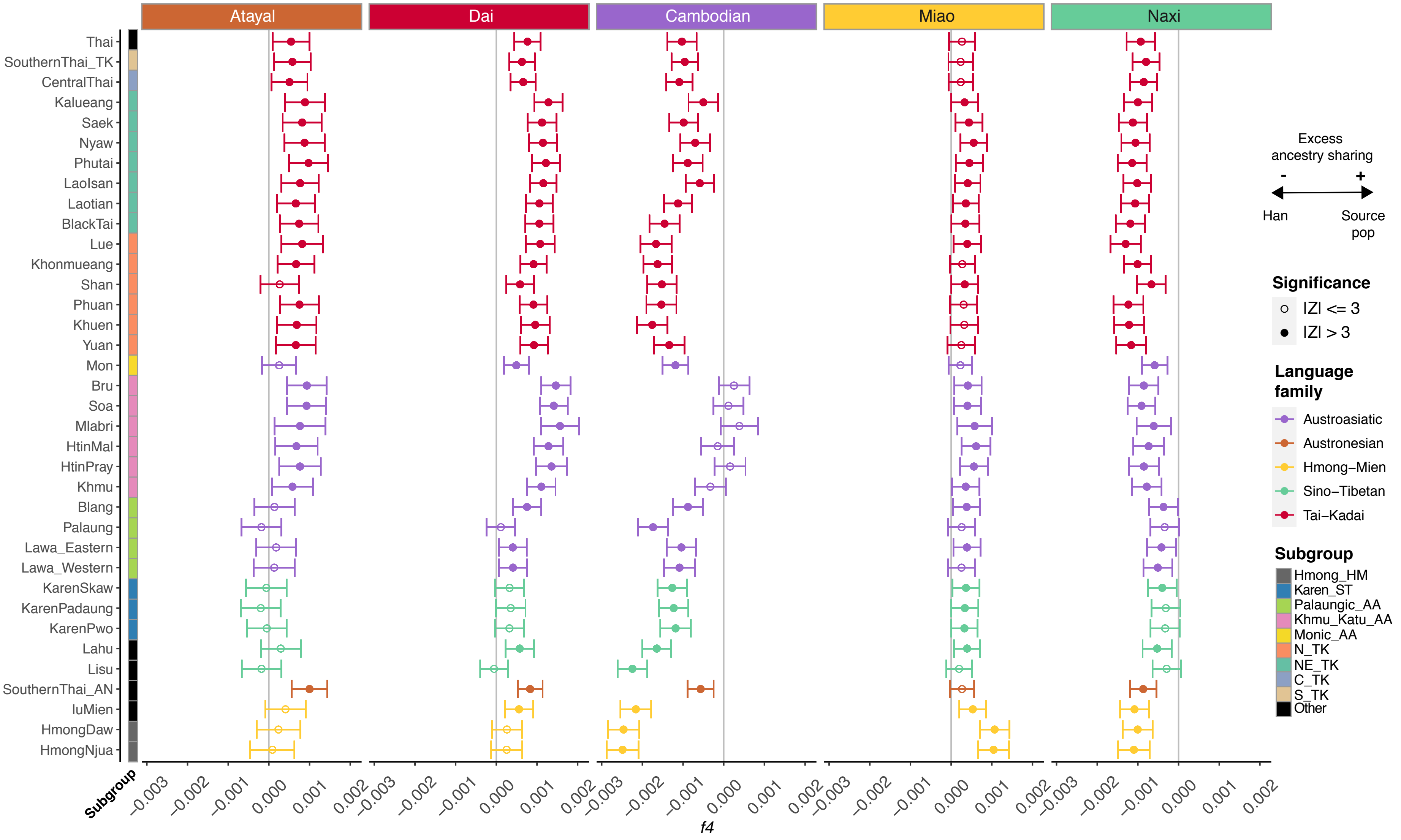


**Supplementary Figure 12** *f4* statistics comparing Thai/Lao populations to representative East Asian populations and Han Chinese. Z-scores are for *f4* (W, Han; Y, Mbuti), where W is the selected East Asian population (panel labels) and Y is the Thai/Lao population (label on the Y axis). The vertical grey lines denote 0. The panels, dots, and error bars are colored according to language family. Empty circles denote nonsignificant Z-scores (|Z|=<3) and solid circles denote significant Z-scores (|Z|>3). The colored bar denotes subgroups of Thai/Lao groups, according to the key on the right.

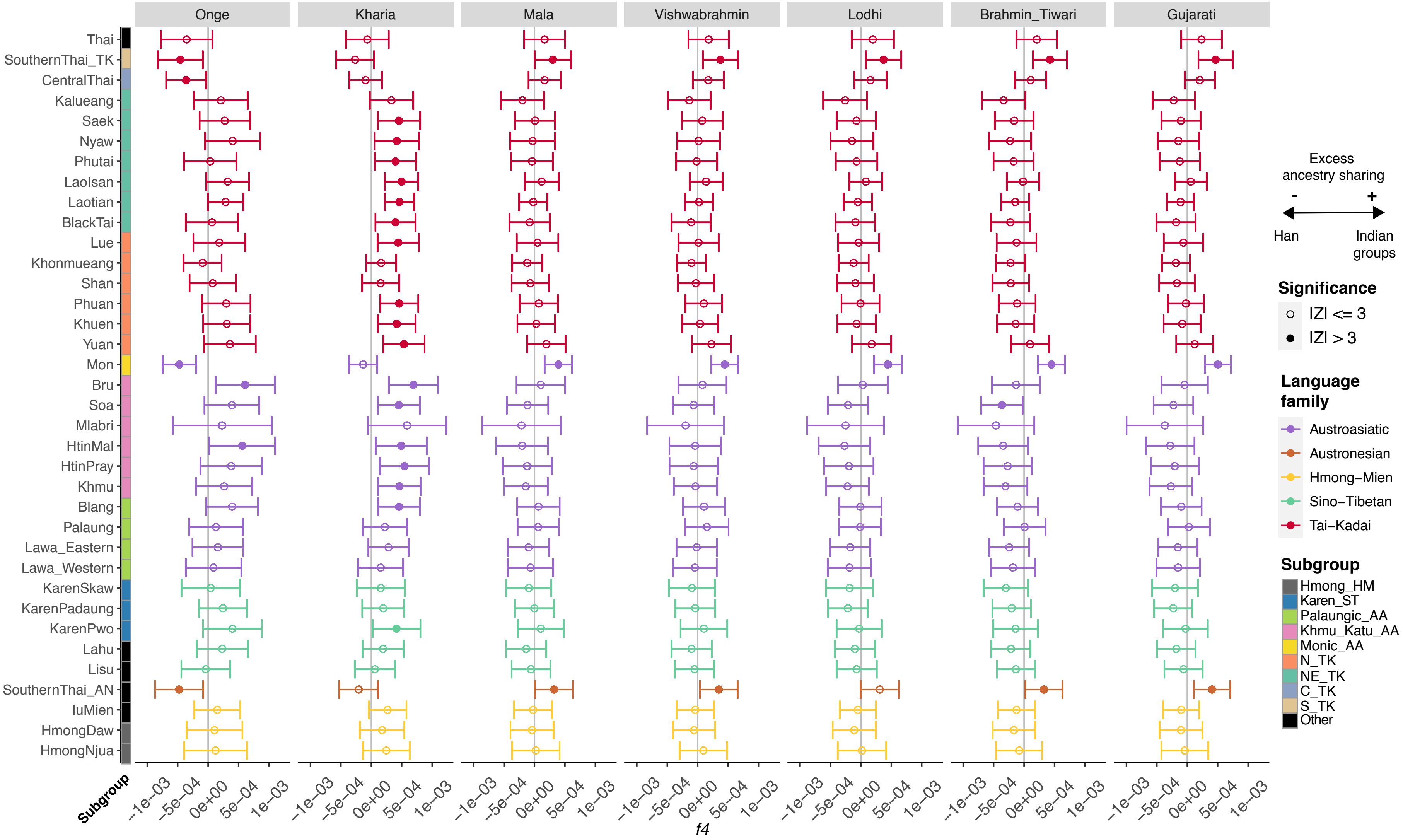


**Supplementary Figure 13** *f4* statistics comparing Thai/Lao populations to Indian populations. Z-scores are for *f4* (Thai/Lao population, Han; Indian population, Mbuti); each panel shows the results for the Indian population (panel label) compared to all Thai/Lao groups (Y-axis labels). The vertical grey lines denote 0. The dots, and error bars are colored according to language family. Empty circles denote nonsignificant Z-scores (|Z|=<3) and solid circles denote significant Z-scores (|Z|>3). The colored bar to the right of the group names denotes linguistic/geographic subgroups of Thai/Lao groups, according to the key on the right.

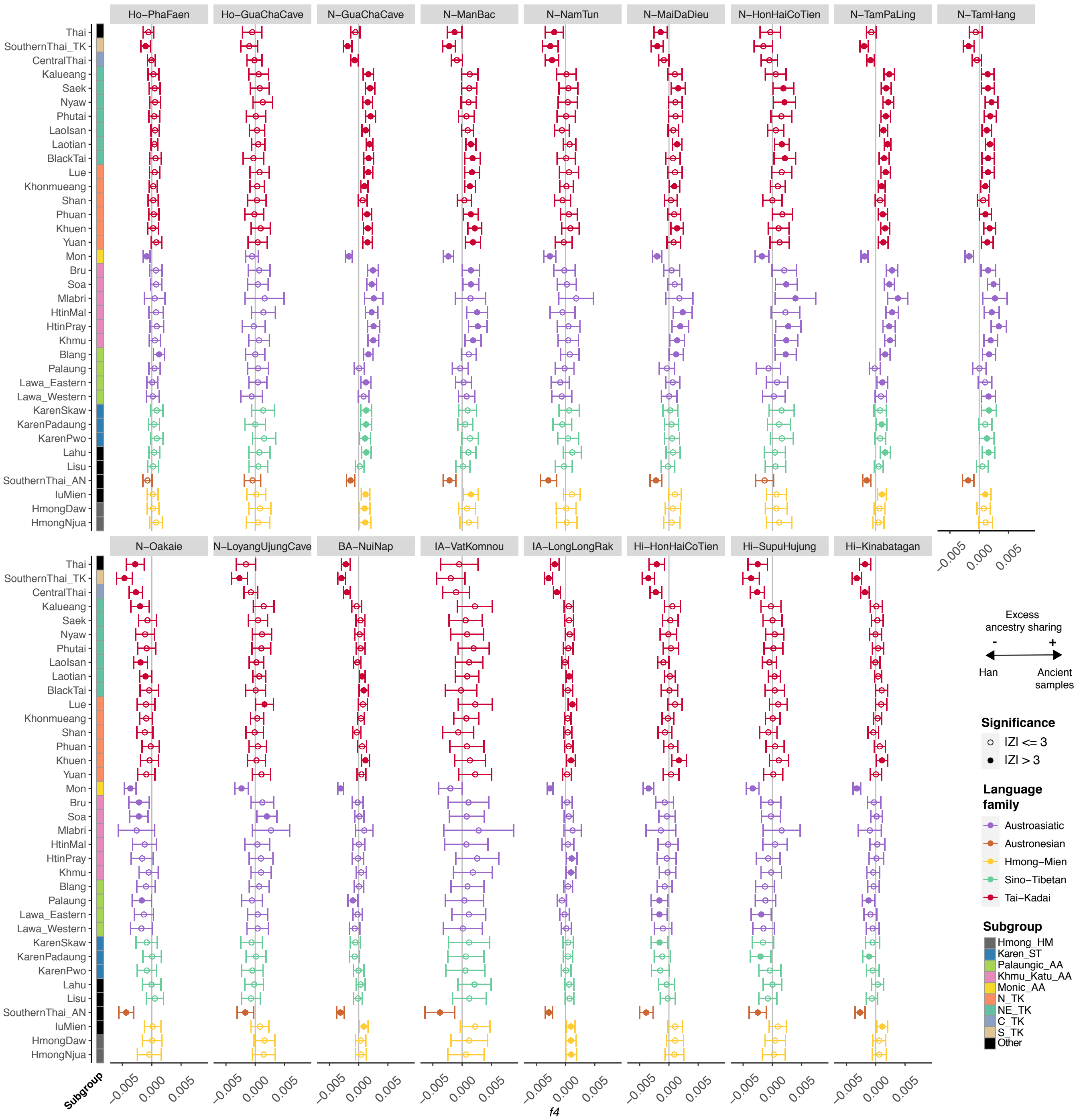


**Supplementary Figure 14** *f4* statistics comparing Thai/Lao populations to ancient samples. Z-scores are for f4 (ancient samples, Han; Thai/Lao populations, French). The vertical grey lines denote 0. The dots and error bars are colored according to language family. Empty circles denote nonsignificant Z-scores (|Z|=<3) and solid circles denote significant Z-scores (|Z|>3). The colored bar denotes subgroups of Thai/Lao groups, according to the key on the right.

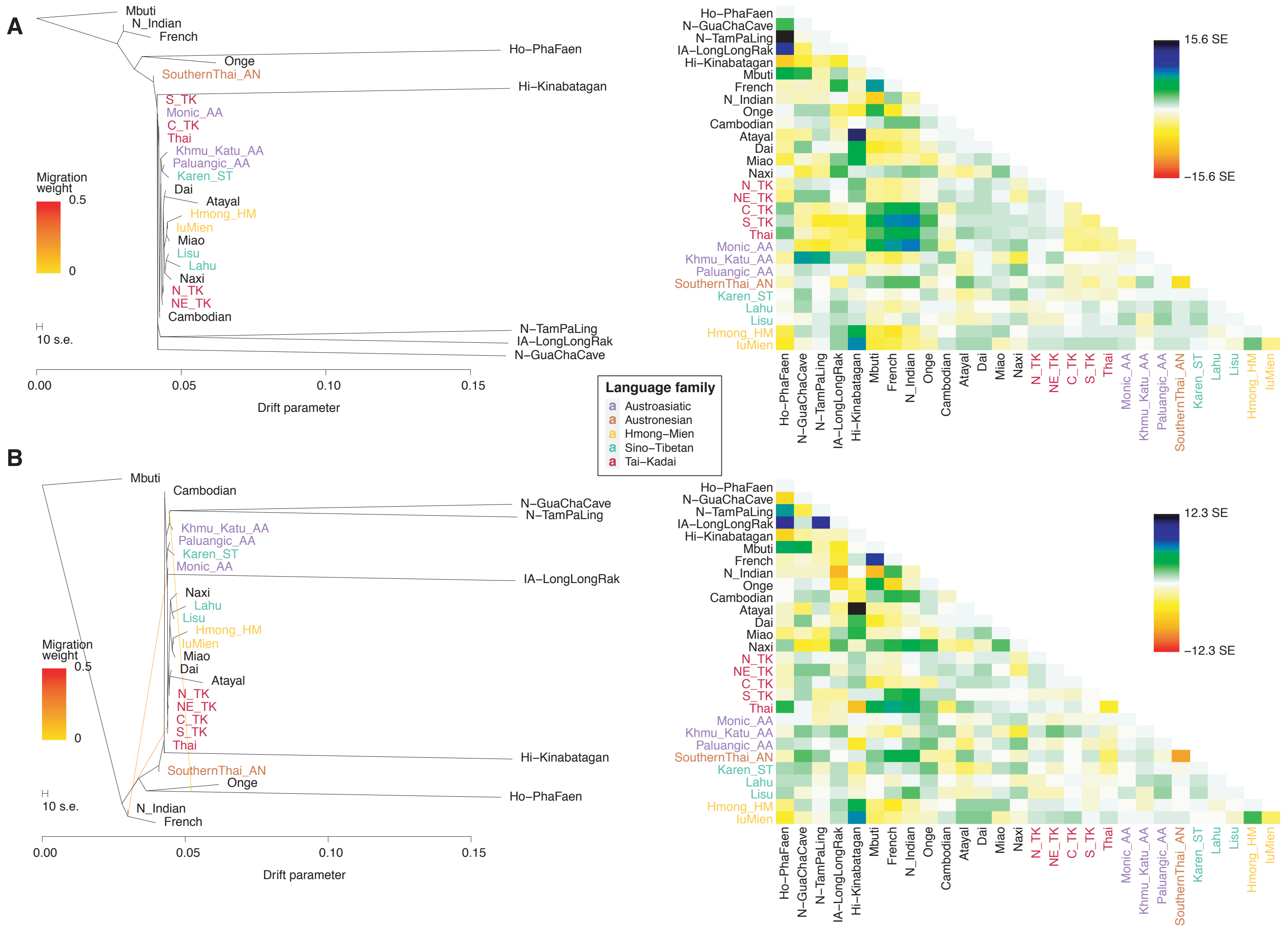


**Supplementary Figure 15** TreeMix results for the Thai/Lao groups and representative modern populations and ancient samples, (A) The maximum likelihood tree without migration and the corresponding residual plot and (B) the maximum likelihood tree with three migration events and the corresponding residual plot. Thai/Lao groups are colored according to language family.

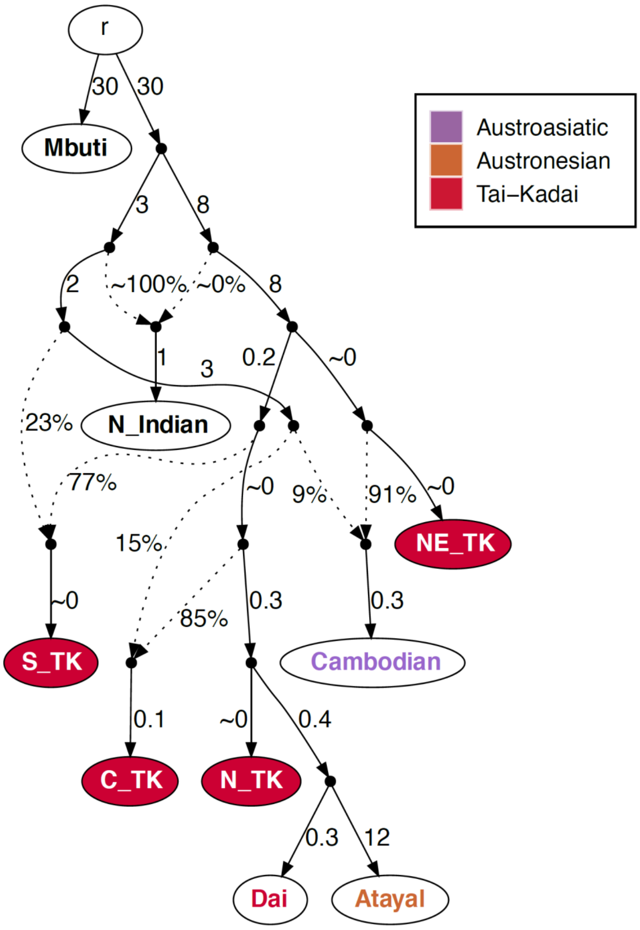


**Supplementary Figure 16** Alternative admixture graph for the TK groups with EA source populations included (worst-fitting Z = -7.037). The node r denotes the root. White nodes denote backbone populations. Backbone population labels and Thai nodes are colored according to language family. Dashed arrows represent admixture edges, while solid arrows are drift edges reported in units of FST×1,000.

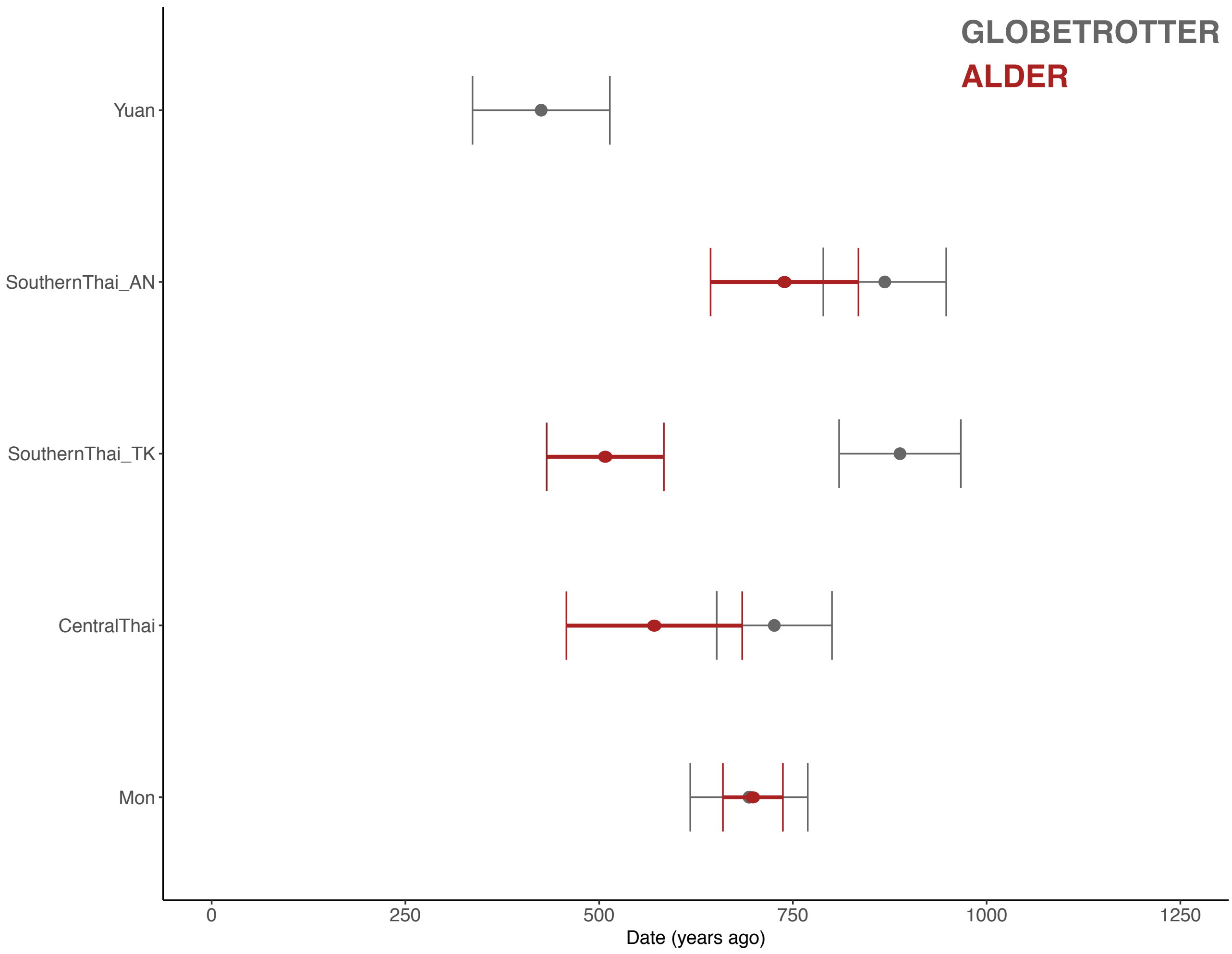


**Supplementary Figure 17** Estimated dates for the SA-related admixture in four putative SA-influenced Thai groups (SouthernThai_AN, SouthernThai_TK, CentralThai and Mon). ALDER estimated dates are in red, using Gujarati as a single source; GLOBETROTTER estimated dates are in grey. Note that ALDER dating failed for the Yuan.

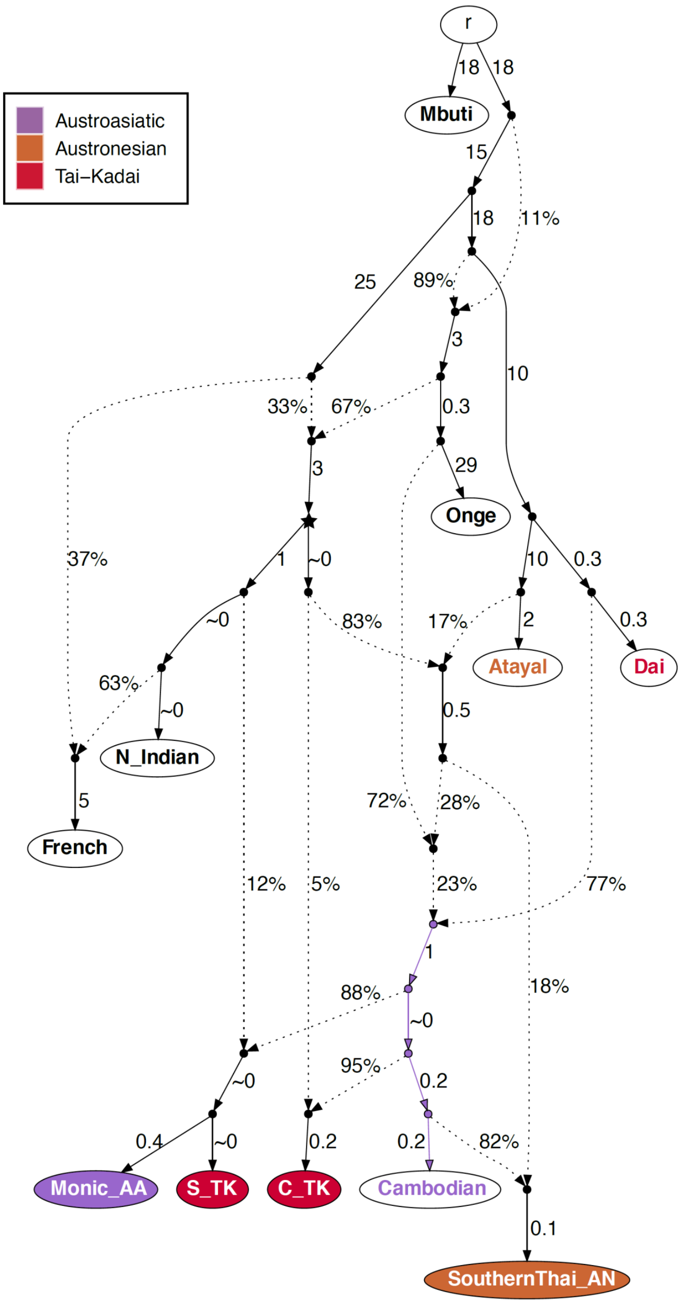


**Supplementary Figure 18** Alternative admixture graph for the SA-related Thai groups with EA source populations included and Mbuti as outgroup (worst-fitting Z = -4.110). The node r denotes the root. White nodes denote backbone populations. Backbone population labels and Thai nodes are colored according to language family. Dashed arrows represent admixture edges, while solid arrows are drift edges reported in units of FST×1,000. The star-shaped node denotes the N_Indian-related source contributing to all of the SA-related Thai groups. The AA-related lineage is highlighted in purple.
